# Evolutionary analysis of the LORELEI gene family in angiosperms reveals regulatory subfunctionalization

**DOI:** 10.1101/2020.04.27.062893

**Authors:** Jennifer A. Noble, Nicholas V. Bielski, Ming-Che James Liu, Thomas A. DeFalco, Martin Stegmann, Andrew D.L. Nelson, Kara McNamara, Brooke Sullivan, Khanhlinh K. Dinh, Nicholas Khuu, Sarah Hancock, Shin-Han Shiu, Cyril Zipfel, Alice Y. Cheung, Mark A. Beilstein, Ravishankar Palanivelu

**Author notes:** **Author Contributions** JAN, NVB, MJL, TAD, MS, KM, BS, KD, NK, and SH planned and designed the research in consultation with and guidance of their respective principal investigators, performed experiments, collected and/or analyzed data. SS performed the transcription factor binding site analysis in the promoters of *LLG* family members. JAN, NVB, ADLN, and MAB performed the evolutionary and comparative transcriptome analyses. JAN, MJL, NVB, TAD, MS, ADLN, SS, CZ, AYC, MAB, and RP wrote the manuscript with input from and revisions by all authors. writing. RP and MAB agree to serve as the authors responsible for contact and communication.

## Abstract

A signaling complex comprising members of the LORELEI (LRE)-LIKE GPI-anchored protein (LLG) and *Catharanthus roseus* RECEPTOR-LIKE KINASE 1-LIKE (*Cr*RLK1L) families perceive RAPID ALKALINIZATION FACTOR (RALF) peptides and regulate growth, reproduction, immunity, and stress responses in Arabidopsis. Genes encoding these proteins are members of multi-gene families in most angiosperms and could generate thousands of signaling complex variants. However, the link(s) between expansion of these gene families and the functional diversification of this critical signaling complex as well as the evolutionary factors underlying the maintenance of gene duplicates remain unknown. Here, we investigated *LLG* gene family evolution, function, and expression in angiosperms. We found that *LLGs* in monocots and eudicots are descendants of a duplication early in angiosperm evolution and that both ancient and recent *LLG* duplicates are retained. Complementation and expression analysis showed that expression divergence of *LLGs* (regulatory subfunctionalization), rather than functional divergence, explains the retention of paralogs in Brassicales. All but one extant monocot and eudicot species examined maintained an *LLG* copy with preferential expression in male reproductive tissues, with the other duplicate copies showed highest levels of expression in female or vegetative tissues. Interestingly, the single *LLG* copy in Amborella (sister to all other angiosperms) is expressed vastly higher in male compared to female reproductive or vegetative tissues. Reconstruction of expression evolution showed that the highest inferred expression levels for the single copy ancestral angiosperm *LLG* was in male reproductive tissues. We propose that expression divergence played an important role in maintenance of *LLG* duplicates in angiosperms.

**One Sentence Summary:** Expression divergence played an important role in maintenance of two sub-groups of *LLG* duplicates in angiosperms

## Introduction

How cells communicate with each other is a fundamental question in eukaryotes. In *Arabidopsis thaliana* (hereafter, Arabidopsis), multimeric signaling complexes comprising members of *Catharanthus roseus* RECEPTOR-LIKE KINASE 1-LIKE (*Cr*RLK1L), LORELEI (LRE)-LIKE GPI-anchored protein (LLG), and RAPID ALKALINIZATION FACTOR (RALF) families influence plant fitness by regulating cell-cell communication during growth, reproduction, immunity, and stress responses (Johnson et al., 2019). *Cr*RLK1Ls are receptor kinases (RKs), LLGs are membrane-associated glycosylphosphatidylinositol-anchored proteins (GPI-Aps), and RALFs are secreted cysteine-rich signaling peptides (CRP). In Arabidopsis, the families that make up *Cr*RLK1L–LLG–RALF signaling complexes are represented by 17, 4, and 37 genes, respectively (Campbell and Turner, 2017; Abarca et al., 2021). Thus, thousands of unique combinations of such a complex may function in diverse processes, distinct tissues, and at different developmental time points. Furthermore, the extent to which such functions in Arabidopsis are conserved in other plants remains unknown (Mecchia et al., 2020). Evolutionary analysis-guided functional studies of multimeric signaling complex members offer unique opportunities to answer this question and dissect the impact of signaling at the cellular level on traits underlying fitness.

Arabidopsis *LRE* is primarily expressed in the synergid cells of the female gametophyte located within the ovules, and plays a role in reproduction (Johnson et al., 2019). Although a potential RALF ligand in this context is still unknown, LRE interacts with three C*r*RLK1Ls - FERONIA (FER), HERCULES RECEPTOR KINASE 1 (HERK1), and ANJEA (ANJ) - to form complexes at the interface of the synergid cell and the incoming pollen tube, rupture the pollen tube to release sperm cells, and effect double fertilization (Duan et al., 2014; Li et al., 2015; Liu et al., 2016; Galindo-Trigo et al., 2020). Consequently, failure to induce pollen tube reception in *fer* or *lre* single mutant pistils or in *herk1 anj* double mutant pistils, leads to ∼80% reduction in seed set (Escobar-Restrepo et al., 2007; Liu et al., 2016; Galindo-Trigo et al., 2020). Elsewhere in the pistil, a variant of this complex (FER-ANJ-HERK1-LLG1-RALF33/RALF23) functions in the stigma and facilitates pollen hydration (Liu C, 2021).

Downstream effects of *Cr*RLK1L-LRE signaling complex have been characterized. LLG1 (in stigma) or LRE (in synergids) and FER initiate a Rho GTPase-mediated signaling pathway that promotes reactive oxygen species (ROS) production via NADPH oxidases of the RESPIRATORY BURST OXIDASE HOMOLOG (RBOH) family that is critical for pollen hydration in stigma and pollen tube reception in synergids (Duan et al., 2014; Liu C, 2021). After pollen tube arrival in the syngergids, FER mediates three key events in the ovule: changes in the synergid calcium profiles (Ngo et al., 2014), relocalization of NORTIA (NTA) from synergid cytoplasm to the filiform apparatus (Yuan *et al*., 2020), and accumulation of nitric oxide in the filiform apparatus to prevent polyspermy (Duan et al., 2020).

In Arabidopsis male reproductive tissues, a variant of this multimeric signaling complex involving the *Cr*RLK1Ls ANXUR1 (ANX1), ANX2, BUDDHA’S PAPER SEAL1 (BUPS1), and BUPS2, along with RALF4/19 and LLG2/3 controls pollen tube integrity *in vivo* (Ge et al., 2017; Zhu et al., 2018; Feng et al., 2019; Ge et al., 2019). Consistent with their preferential expression in pollen grains and pollen tubes, loss-of-function mutants in members of this complex result in swollen tips, and/or premature pollen tube burst soon after germination (Feng et al., 2019; Ge et al., 2019). Additionally, RNAi mutant pollen tubes, in which both *LLG2* and *LLG3* are expressed at low levels, exhibited decreased pollen tube length *in vitro* and *in vivo* (Feng et al., 2019; Ge et al., 2019). Slowed pollen tube growth in RNAi *llg2 llg3* pollen tubes likely resulted from insufficient secretion of cell wall components, which is linked to increased accumulation of esterified and de-esterified pectin in the apical and subapical region, respectively, and decreased accumulation of callose (Feng et al., 2019). Upon activation, the ANX-based signaling pathway, coupled with pollen tube-expressed RBOHs, is important for ROS production and for maintenance of a steady tip-focused Ca^2+^ gradient that sustains pollen tube growth (Boisson-Dernier et al., 2013).

Besides reproduction, variants of the *Cr*RLK1L–LLG–RALF signaling complex mediate diverse processes in vegetative tissues, stress responses, and immunity in Arabidopsis (Duan et al., 2010; Li et al., 2015; Stegmann et al., 2017; Feng et al., 2018; Xiao et al., 2019) and the crystal structure of the ternary FER-LLG2-RALF23 complex has been solved (Xiao et al., 2019). The FER–LLG1–RALF1 complex signals via a Rho GTPase complex to regulate ROS production and is critical for root hair growth and hypocotyl elongation (Duan et al., 2010; Li et al., 2015). Consequently, *fer* and *llg1* mutants show similar vegetative defects, which includes reduced rosette size and many root abnormalities, including insensitivity to RALF1-dependent growth inhibition, decreased sensitivity to auxin-stimulated ROS production, sensitivity to abscisic acid, and abnormal root hair development (Li et al., 2015). Although the malectin-like domains (MLDs) of CrRLK1Ls do not present conserved carbohydrate-binding residues, the extracellular domain of FER binds de-esterified pectin and this interaction is likely important for maintaining Arabidopsis root cell integrity when subjected to salt stress and restoring growth recovery after exposure to high salinity (Feng et al., 2018). A similar FER–LLG1–RALF23 complex regulates immunity by modulating the perception of pathogen-associated molecular patterns (PAMPs) such as flagellin (or the corresponding peptide epitope flg22) and elongation factor thermo unstable (EF-TU, or the corresponding peptide epitope elf18). In this context, RALF23 is recognized by its FER–LLG1 receptor complex to suppress the scaffolding function of FER, thereby inhibiting ROS production and immunity (Stegmann et al., 2017).

Other components that could interact with this *Cr*RLK1L–LLG–RALF module to form higher-order signaling complexes include EARLY NODULIN LIKE-14 (EN14) in the synergid cells and LEUCINE-RICH REPEAT EXTENSIN (LRX) proteins in the pollen tubes and vegetative tissues. EN14, which is a GPI-AP with a plastocyanin-like domain, interacts with FER and LRE and loss-of-function mutants in *EN11-EN15* contained ovules with pollen tube reception defects and ∼50% reduction in seed set (Hou et al., 2016). LRX proteins bind RALF peptides and help in maintaining pollen tube integrity (Mecchia et al., 2017; Moussu et al., 2020). LRXs also associate with FER in vegetative tissues, where they regulate immunity (Gronnier et al., 2020), salt stress tolerance (Zhao et al., 2018), as well as root hair growth (Herger et al., 2020) and vacuolar expansion (Dunser et al., 2019).

Phylogenetic analysis of each component of the *Cr*RLK1L–LLG–RALF multimeric complex is essential to elucidate the links between cell-cell communication and the evolution of traits that influence plant fitness. Phylogenetic analysis of 1050 *CrRLK1L* proteins in 57 plants showed that the C*r*RLK1L subfamily has undergone several duplication events during its evolution and that expression of C*r*RLK1Ls in legumes and Arabidopsis were conserved (Solis-Miranda et al., 2020). In another study, the phylogenetic relationship among *Cr*RLK1L genes identified retention of duplicates in Arabidopsis, moss, rice, medicago, and tomato and that their evolution pointed to functional conservation (Yang et al., 2021). Phylogenetic analysis of 795 RALFs from 51 land plants showed that evolution of the RALF family was distinct in eudicots and monocots and that numerous RALF genes in Arabidopsis resulted from a rapid expansion in the Brassicaceae (Campbell and Turner, 2017). However, a comprehensive evolutionary analysis paired with functional studies and expression profiling has not been performed for any of the *Cr*RLK1L–LLG–RALF signaling complex members. Consequently, links between expansion of these gene families and the functional diversification of this critical signaling complex and evolutionary factors underlying the maintenance of gene duplicates remains unknown.

In this study, we addressed maintenance of duplicate members of the *LLG* family by inferring phylogeny and characterizing patterns of functional and regulatory evolution to investigate if ancestral functions are shared between duplicated copies (subfunctionalization) or if the ancestral expression pattern is divided between the duplicates, leading to non-overlapping expression of each paralog (regulatory subfunctionalization) (Ohno, 1970; Force et al., 1999; Des Marais and Rausher, 2008). We identified homologs of the four-member Arabidopsis *LLG* gene family and showed that they are conserved throughout angiosperms. We used complementation assays and found that the molecular functions of GPI-APs encoded by the *LLG* gene family are conserved. In contrast, expression of *LLG* family members in representative angiosperms is highly divergent showing that regulatory divergence (i.e., differences in gene expression), rather than functional divergence, likely contributed to the retention of *LLG* family members in angiosperms. In all but one of the extant monocots and eudicots species, we found that at least one *LLG* copy is preferentially expressed in male reproductive tissues, with the other paralog(s) showing highest levels of expression in female or vegetative tissues. Interestingly, the single *LLG* copy in the angiosperm *Amborella trichopoda* (hereafter, Amborella) is expressed higher in male reproductive tissues compared to female or vegetative tissues. Additionally, likelihood reconstruction of ancestral expression across plant tissues revealed that elevated *LLG* expression in pollen and pollen tube is ancient. Taken, together, our analyses suggest that the *Cr*RLK1L-LLG-RALF signaling complex likely plays a role in pollen tube growth regulation across angiosperms.

## Results

### Maintenance of duplicated LLG gene families is widespread in angiosperms

To understand the evolutionary history of the four *LLGs* in Arabidopsis (Capron *et al*., 2008), we inferred phylogeny for *LLG* homologs recovered from angiosperm genomes. Our motif-driven homology searches (Bailey and Gribskov, 1998) and subsequent filtering returned 114 putative *LLG* family members across 32 focal genomes (Dataset S1). We rooted the tree resulting from our maximum likelihood search with the single *LLG* from Amborella, a putative sister species to all other extant angiosperms (The Angiosperm Phylogeny et al., 2016). The tree indicated that *LLGs* in monocots and eudicots experienced independent duplication events after the two lineages diverged (Figure 1). For this reason, we alphabetized homologs recovered in monocots (*LLGA-H,* Figure 1) to reflect their independent evolution and distinguish them from eudicot *LLGs*, which are numbered. Our tree indicated that *LLGs* in monocots likely underwent several lineage-specific duplication events, including one that resulted in two *LLG* clades in grasses (*LLGA* and *LLGB*) (Figure 1). Importantly, all the monocot genomes included here retained duplicate *LLGs*.

**Figure 1.**
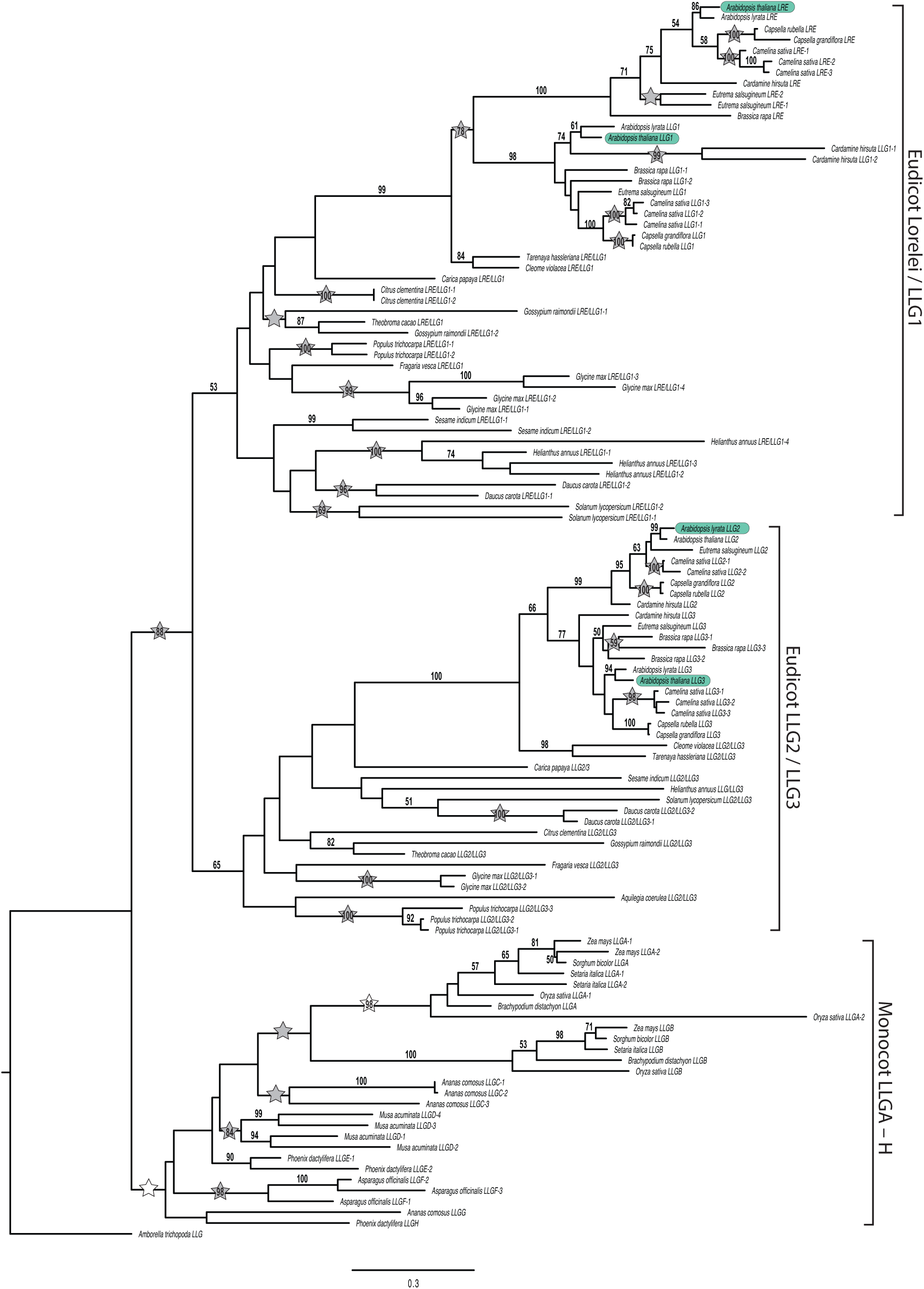
Full LLG family phylogeny across angiosperms. Maximum likelihood tree (ln(L) = -33601.508911) inferred using RAxML (RAxMLHPC MPI AVX2 v. 8.2.11) on a partitioned nucleotide alignment. Support for nodes was assessed using the rapid bootstrap algorithm (1000 replicates). The tree is rooted with the single copy Amborella *LLG*. Grey stars indicate gene duplication events. White stars indicate duplications that may be impacted by phylogenetic uncertainty. Monocots and eudicots experienced independent duplication events. LLGs duplicated early in eudicot evolution resulting in reciprocally monophyletic *LRE/LLG1* and *LLG2/LLG3* clades. In Brassicaceae, a subsequent set of duplication events yielded the four copies present in *Arabidopsis* (indicated in green).

Eudicot *LLG* sequences in our tree descended from a common eudicot ancestor (Figure 1, 88% bootstrap support) that underwent an early duplication event yielding the *LRE/LLG1* and *LLG2/LLG3* copies present in extant eudicots. Except for *Aquilegia coerulea*, which retains only an *LLG2/LLG3* ortholog, sampled eudicots retained both descendants of this early duplication. In Brassicaceae, a subsequent family-wide duplication and retention yielded four *LLGs*. In Brassicaceae, *LRE* and its orthologs formed a monophyletic group sister to the clade containing all *LLG1* orthologs. Similarly, *LLG2* and its orthologs formed a monophyletic group sister to the clade containing all *LLG3* orthologs (Figure 1). In sum, the evolutionary history of the *LLG* family is characterized by the retention of both ancient and recent duplicates in angiosperms.

### LLG molecular functions are conserved in Arabidopsis and members of Cleomaceae

The retention of duplicate genes typically results from evolutionary mechanisms that can produce signatures of selection and divergence in promoter regions, coding sequence, or both (Force et al., 1999; Li et al., 2005; Duarte et al., 2006). To test if retention of the *LRE/LLG1* and *LLG2/LLG3* copies present in extant eudicots resulted from changes in coding sequence thereby yielding distinct functions after an early duplication event in their evolution, we examined *LLG*s in Arabidopsis, where characterized mutants made functional analysis feasible (Li et al., 2015; Liu et al., 2016; Ge et al., 2019). We expressed all Arabidopsis *LLG* coding sequences from the Arabidopsis *LLG1* promoter and tested if they could complement the vegetative development defects of *llg1-2* mutants (Li et al., 2015). Like *LLG1,* expression of Arabidopsis *LRE* or *LLG3* from the *LLG1* promoter complemented root hair defects (Figure 2a-d), hypocotyl length in dark-grown *llg1-2* seedlings (Figure 2e), epidermal pavement cell defects (Figures 3a-3e), and RALF1 sensitivity in *llg1-2* seedlings (Figures 3g and S1).

**Figure 2.**
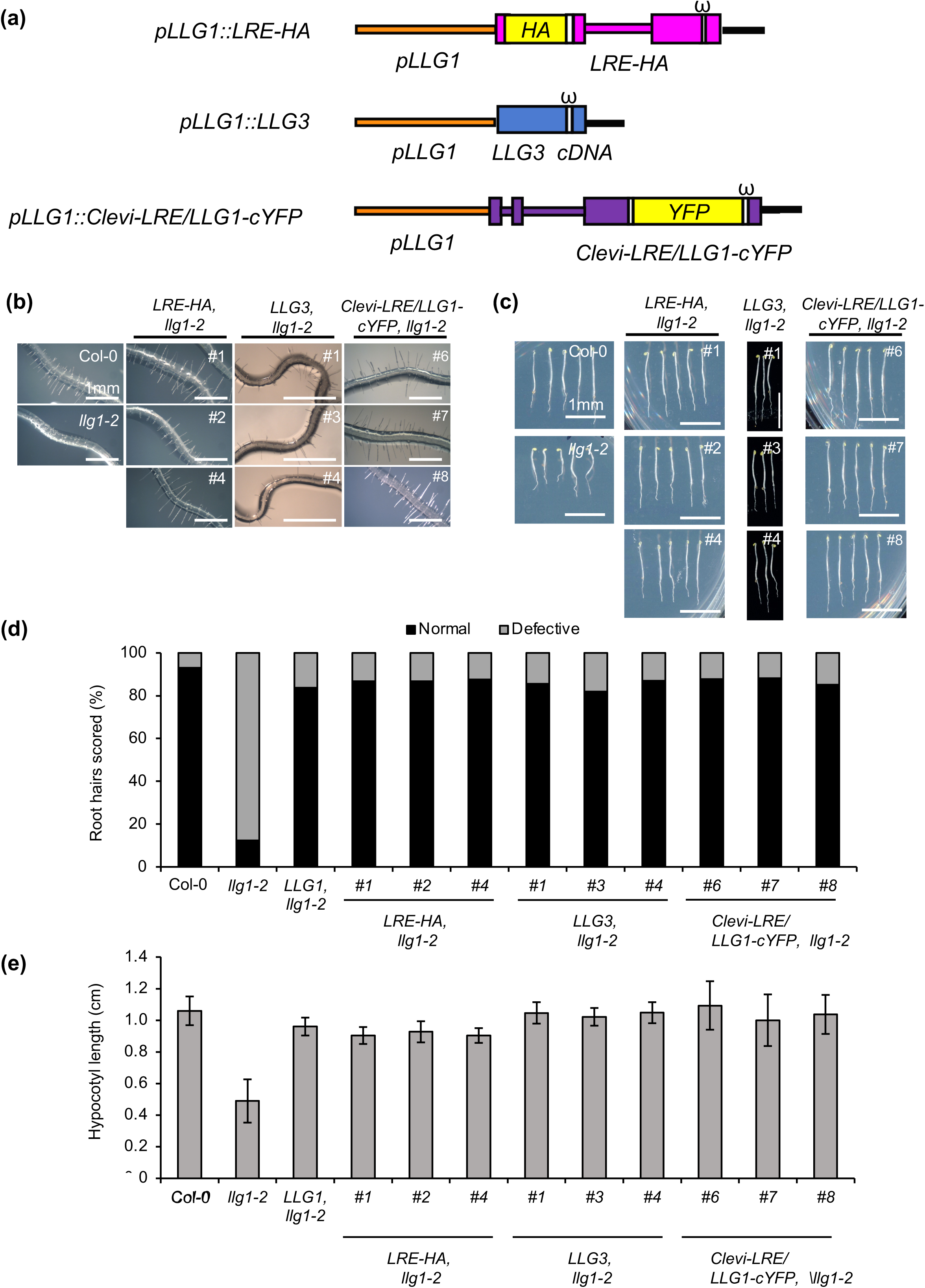
LRE-HA, LLG3, and Clevi-LRE/LLG1-cYFP complemented vegetative defects in *llg1-2* seedlings. (a) Diagrams of the *pLLG1::LRE-HA, pLLG1::LLG3,* and *pLLG1::Clevi-LRE/LLG1-cYFP* constructs. (b) Root hair length defects were restored when LRE-HA, LLG3, and Clevi-LRE/LLG1-cYFP were expressed in *llg1-2* seedlings to wild-type (Col-0) levels. Images were taken from T2 seedlings of single insertion lines. (c) LRE-HA, LLG3, and Clevi-LRE/LLG1-cYFP complementation of short hypocotyl length phenotype in *llg1-2* seedlings. Representative images of wild-type (Col-0) or T2 seedlings of single-locus insertion lines of *pLLG1::LRE-HA, pLLG1::LLG3,* or *pLLG1::Clevi-LRE/LLG1-cYFP* are shown. (d) Quantification of complementation of root hair defects in *llg1-2* seedlings shown in B are reported here. Root hairs were scored as normal or defective in at least 10 seedlings in each indicated single-insertion line carrying LRE-HA, LLG3, or Clevi-LRE/LLG1-cYFP fusion protein. (e) Quantification of complementation of de-etiolated hypocotyl in dark grown *llg1-2* mutant seedlings shown in C are reported here. Hypocotyl length was assayed in T2 seedlings from selfed seeds of single insertion lines carrying LRE-HA, LLG3, or Clevi-LRE/LLG1-cYFP fusion protein. Quantification of hypocotyl lengths was done in at least three trials of 20-25 seedlings for each line. Error bars represent ±SD.

**Figure 3.**
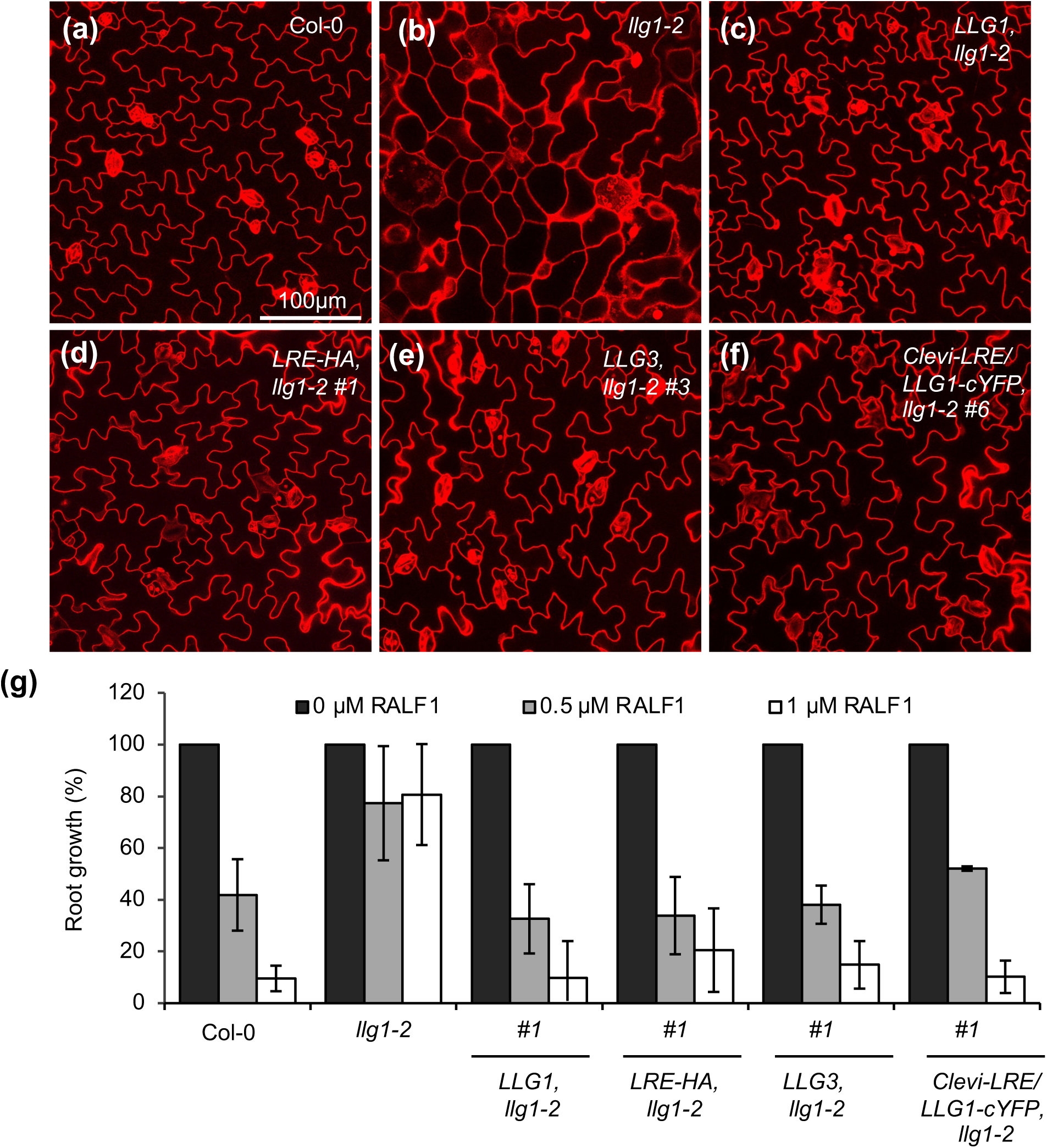
LRE-HA, LLG3, and Clevi-LRE/LLG1-cYFP complemented epidermal defects and RALF1 insensitivity in *llg1-2* seedlings. (a-f) Epidermal pavement cells of 6-day-old *llg1-2* seedling cotyledons expressing LRE-HA, LLG3, or Clevi-LRE/LLG1-cYFP showed restored normal pavement cell morphology like that seen in wild-type (Col-0) and *llg1-2* seedlings expressing LLG1. In each genotype, 5-10 seedlings were stained with Propidium Iodide (PI) and visualized with confocal microscopy. (g) Percentage of root growth after RALF1 treatment of three-day-old seedlings as described in Figure S2 Root length was measured two days after RALF1 treatment, and three trials were performed. Error bars represent ±SD.

Expression of Arabidopsis *LLG2* also complemented vegetative defects in *llg1-2* mutants, as rosette size was restored to levels observed with *LLG1* (Figure 4a). In addition to vegetative phenotypes, *llg1* mutants are defective in immune responses, as they show reduced responsiveness to several PAMPs, including the bacterial elicitors elf18 and flg22 (Shen et al., 2017; Stegmann et al., 2017; Xiao et al., 2019). In *llg1-2* plants expressing *LLG2* from the *LLG1* promoter, we found that responses to elf18 and flg22 were restored to levels comparable to those expressing *LLG1* (Figures 4b and S2). Similarly, complementation of *llg1-2* with *LLG2* restored responsiveness to exogenous RALF23 peptide to levels detected in *llg1-2* plants expressing *LLG1* (Figure 4c). These results demonstrated that LRE, LLG2, or LLG3 can substitute for LLG1 and perform its molecular functions. Results of these complementation experiments are particularly significant, as *LLG2* and *LLG3* are members of a lineage that is distinct from *LLG1* and originated from an ancient duplication event early in the eudicot evolution (Figure1).

**Figure 4.**
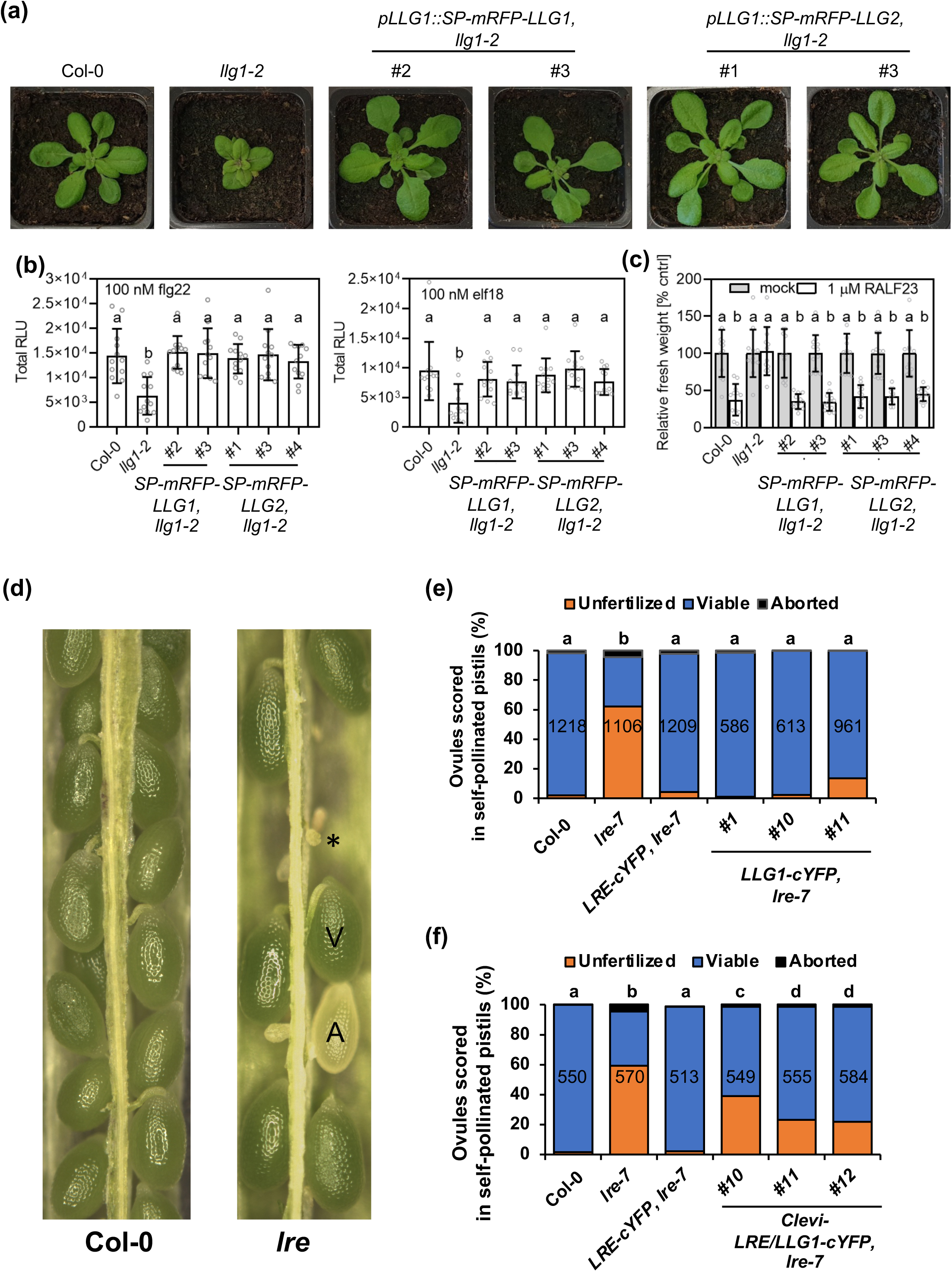
mRFP-LLG1 and mRFP-LLG2 complemented defects in *llg1-2* and LLG1-cYFP and Clevi-LRE/LLG1-cYFP complemented reproductive defects in *lre.* (a) Expression of *pLLG1::SP-mRFP-LLG*1 or *pLLG1::SP-mRFP-LLG2* restores rosette size in 4.5-week-old *llg1-2* plants. (b) ROS production in response to flg22 (left) or elf18 (right) is restored in SP-mRFP-LLG1 or SP-mRFP-LLG2 lines driven by the *LLG1* native promoter. Total Relative Light Unit (RLU) over 40 minutes of exposure to 100 nM flg22 or elf18 treatment is displayed. Letters indicate significantly different values (n=12 leaf discs, two-way ANOVA with Tukey test, flg22 p<0.0001; elf18 p=0.0009). Error bars show ±SD. (c) Growth sensitivity to RALF23 is restored in seedlings expressing SP-mRFP-LLG1 or SP-mRFP-LLG2. Letters indicate significantly different values (n=16 seedlings, two-way ANOVA with Tukey test, p<0.0001). Error bars show ±SD. (d) Images of opened siliques of indicated genotypes in *Arabidopsis.* A representative unfertilized ovule (*) and viable (V) or aborted (A) seed is marked in the *lre* silique. (e) LLG1-cYFP complemented *lre* mutant seed set defects in self-pollinated pistils of indicated three independent transformants (ANOVA, p = 0.18). (f) Clevi-LRE/LLG1-cYFP partially complemented *lre* mutant seed set defects in self-pollinated pistils of indicated three independent transformants (pairwise two-tailed t-tests, p > 0.05). (e, f) Number in the middle of each column refers to the number of ovules/seeds scored. Groups sharing the same lowercase letters are like seach other in statistical tests.

We also examined if molecular functions of *LRE* in female reproductive tissues are conserved. For this, we expressed Arabidopsis *LLG1* from the promoter of its closest paralog (Arabidopsis *LRE*) and tested if it could complement the female reproductive defects in the *lre* mutant (Liu et al., 2016). Like LRE fused to citrine YFP (LRE-cYFP) (Liu et al., 2016), LLG1-cYFP localized in the synergid cells (Figure S3) and restored seed set defects in *lre-7* to levels detected in wild type or LRE-cYFP plants based on fertility (Figures 4d and 4e) and transmission assays (Table S1).

To examine conservation of functions in LLGs in other plants besides those in Brassicaceae, we explored *LLGs* in the sister family of Cleomaceae. Unlike Brassicaceae, there are only two single copy orthologs in the Cleomaceae – *LRE/LLG1* and *LLG2/LLG3* (Figure 1). Since LRE and LLG1 can reciprocally complement each of their functions in the vegetative and reproductive tissues of Arabidopsis, respectively, we tested if the molecular functions performed by Arabidopsis LRE and LLG1 are conserved in the LRE/LLG1 single copy ortholog from *Cleome violacea* (Clevi-LRE/LLG1). We expressed *Clevi-LRE/LLG1-cYFP* from the Arabidopsis *LLG1* or *LRE* promoter (Figure 2f) in the *llg1-2* or *lre-7* mutant, respectively, and performed complementation experiments. We found that Clevi-LRE/LLG1 complemented root hair defects (Figures 2b and 2d), hypocotyl lengths in dark-grown *llg1-2* seedlings (Figures 2c and 2e), epidermal cell defects (Figure 3f), and RALF1 sensitivity in *llg1-2* seedlings (Figures 3g and S1) to levels seen in *llg-1-2* lines expressing LLG1. Additionally, Clevi-LRE-cYFP expressed from the *LRE* promoter localized in the synergid cells (Figure S3) and partially restored female reproductive defects in *lre-7* based on fertility (Figure 4f) and transmission assays (Table S2). These results indicated that the *C. violacea* single-copy LRE/LLG1 ortholog complements the functions of both Arabidopsis LRE and LLG1. Results from complementation experiments led us to conclude that the molecular functions of LLGs are highly conserved and thus changes to the coding regions of these genes (LRE shares 58%, 55%, and 46% amino acid identity with LLG1, LLG3, and LLG2, respectively (Capron et al., 2008), were likely not responsible for the post-duplication retention of the eudicot copies.

### LLG gene expression is divergent in Arabidopsis and members of Cleomaceae

We next considered if expression divergence among *LLG* paralogs underlies their retention. Consistent with this possibility, members of the *LLG* gene family in Arabidopsis are differentially expressed (Tsukamoto et al., 2010; Feng et al., 2019; Ge et al., 2019; Xiao et al., 2019). *LLG2* and *LLG3* are primarily expressed in male reproductive tissues (Feng et al., 2019; Ge et al., 2019; Xiao et al., 2019) while *LRE* is primarily expressed in female reproductive tissues and seedlings (Tsukamoto et al., 2010; Wang et al., 2017). Unlike the preferential expression of these three *LLGs* in reproductive tissues, *LLG1* is ubiquitously expressed throughout plant development (Tsukamoto et al., 2010; Feng et al., 2019; Ge et al., 2019; Xiao et al., 2019). However, based on RT-PCR and RNA-seq experiments, there is an overlap of *LRE* and *LLG1* expression in seedlings and unpollinated and pollinated ovaries (Tsukamoto et al., 2010; Ge et al., 2019). To test if expression of these two genes overlapped in cells of these complex tissues, we examined cell-specific β ucuronidase (GUS) expression in three *pAtLRE::GUS* lines (this study), and one previously reported *pAtLLG1::GUS* line (Li et al., 2015). GUS expression analyses revealed no cell type in seedlings, unpollinated pistils, or pollinated pistils where *LRE* and *LLG1* are co-expressed (Figure 5), demonstrating that *LLG1* and *LRE* indeed have different and non-overlapping expression profiles.

**Figure 5.**
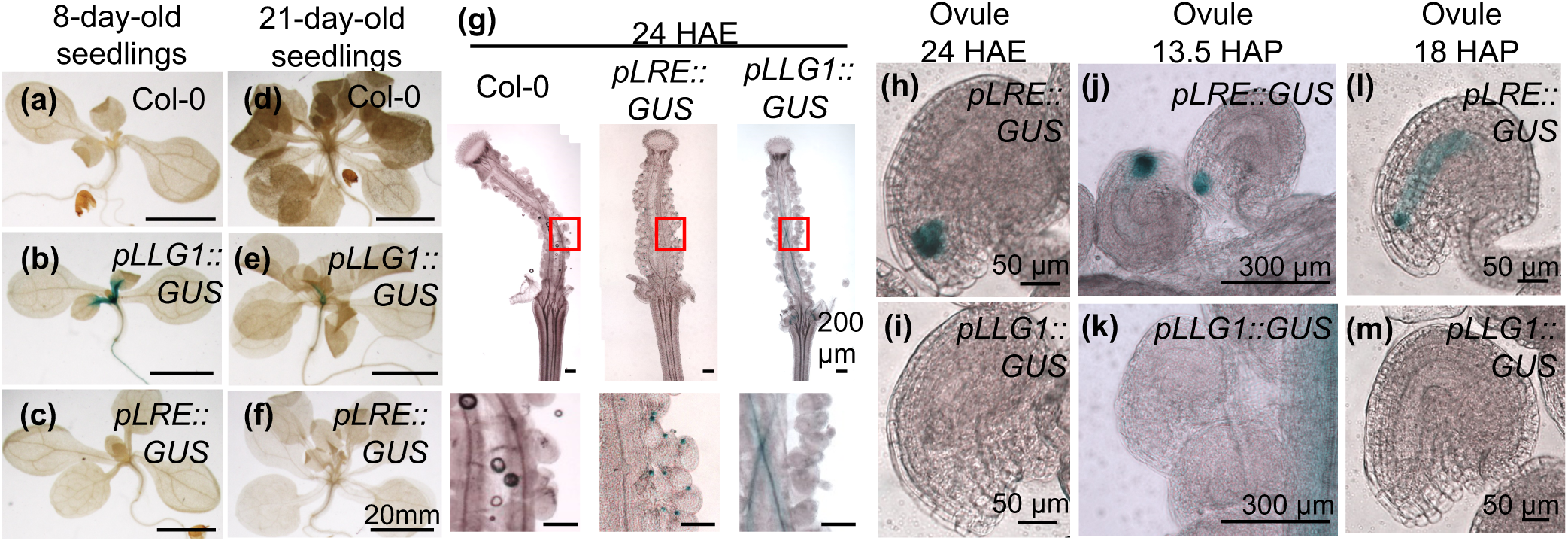
*pLRE::GUS* and *pLLG1::GUS* showed non-overlapping expression in vegetative and reproductive tissues. (a-f) *pLLG1::GUS* was expressed, while *pLRE::GUS* was not, in vegetative tissues. In 8-day-old seedlings, *pLLG1::GUS* was expressed in true leaves, hypocotyls, and roots (a-c). In 21-day-old seedlings, *pLLG1::GUS* was expressed in the epicotyl, the hypocotyl, and weakly expressed in roots (d-f). (g) At 24 hours after emasculation (HAE), *pLRE::GUS* and *pLLG1::GUS* were both expressed in pistils but in different cell-types. *pLRE::GUS* was expressed in synergid cells, while *pLLG1::GUS* was expressed in septum. Close up of the area marked in red rectangles are shown below. (h-i) *pLRE::GUS* was expressed in synergid cells at 24 HAE (h) but *pLLG1::GUS* is not expressed in the ovule (i). (j-m) *pLRE::GUS* and *pLLG1::GUS* showed non-overlapping expression after pollination. Mature unpollinated pistils were pollinated with Col-0 pollen and collected at 13.5 hours after pollination (HAP) (j-k) or 18 HAP (l-m) and stained for GUS activity. (j-k) At 13.5 HAP, *pLRE::GUS* was expressed in the micropylar end of the female gametophyte (j), while *pLLG1::GUS* continues to be expressed in the septum (k). (l-m) At 18 HAP, *pLRE::GUS* and *pLLG1::GUS* were both expressed in pollinated pistils but in different cell-types. *pLRE::GUS* was expressed in the zygote and developing endosperm nuclei, while *pLLG1::GUS* was expressed in septum.

We also explored *LLG* expression divergence in plants other than Arabidopsis by examining published RNA-seq based transcriptome atlases of species in Cleomaceae. Indeed, we found that the single copy *LRE/LLG1* orthologs in *Tarenaya hassleriana* and *Gynandropsis gynandra* were expressed in all developmental stages (Kulahoglu et al., 2014) and encompassed all the expression domains of Arabidopsis *LRE* + *LLG1* (Table S3). In contrast, the single copy *LLG2/LLG3* ortholog in both plants was expressed nearly exclusively in male reproductive tissues (Table S3). These results showed that *LLG* gene expression is divergent in Arabidopsis and members of Cleomaceae, in striking contrast with the conservation of functions among *LLGs*.

### Regulatory subfunctionalization may underlie retention of duplicated LLGs in angiosperms

To examine how widespread expression divergence is among *LLG* family members in other angiosperms, we analyzed published RNA-seq-based transcriptome data (see Methods) and calculated the normalized expression levels of *LLGs* across five expression domains (male and female reproductive tissues, vegetative tissues, and sporophytic reproductive tissues before and after fertilization) (Table S4). We analyzed expression in ten angiosperms for which we recovered *LLG* homologs – four eudicots (Arabidopsis*, T. hassleriana*, *Solanum lycopersicum*, and *Sesamum indicum*) and six monocots (*Oryza sativa, Zea mays, Sorghum bicolor, Brachypodium distachyon, Phoenix* dactylifera, and *Annanas comosus*). We found that expression patterns have diverged post duplication in both eudicots and monocots (Table S4).

Eudicots are characterized by an early duplication that yielded two reciprocally monophyletic clades of *LLGs*: *LRE/LLG1* and *LLG2/LLG3*. Members of the *LRE/LLG1* clade primarily showed the highest level of expression in female and vegetative tissues, while members of the *LLG2/LLG3* clade are expressed at their highest level and almost exclusively restricted to male reproductive tissues (Table S4). In contrast, expression divergence occurred within the *LRE/LLG1* clade. For example, *Solanum lycopersicum* retains two *LRE/LLG1* paralogs; one is expressed in female reproductive tissues and sporophytic tissues before fertilization, while the other is most highly expressed in vegetative tissues (Table S4).

Like our findings in eudicots, duplication in the sampled grass monocots yielded one copy with the highest expression in male reproductive tissues and another copy that was expressed more broadly. For example, the *LLGB* clade members exhibited male reproductive tissue expression. However, members of the sister clade, the grass specific *LLGA*, underwent additional duplication events and expansion of expression across domains. In *O. sativa*, transcription of *LLGA-1* was at its highest levels in the female domain and reproductive tissues before fertilization, while the domain of highest expression for *LLGA-2* was in vegetative tissues (Table S4). Thus, the pollen- and pollen tube-preferential expression patterns observed for Arabidopsis *LLG2* and *LLG3* appeared to mirror that of *LLG2/3* orthologs in other eudicots and *LLGB* copies in grass monocots (Table S4).

Copies of *LLGs* from non-grass monocots (*P. dactylifera* and *A. comosus*) were derived from a combination of older and more recent duplication events (Figure 1). For *P. dactylifera* the pattern of expression divergence observed in other species holds; *LLGH* is restricted to male reproductive tissues and *LLGE-1 and 2* are more broadly expressed, including in female reproductive and vegetative tissues (Table S4). Thus, our analyses revealed that preferential expression of at least one *LLG* copy in male reproductive tissues has independently evolved in eudicots and in monocots (Table S4). Still, in contrast to all other species, *A. comosus* appears to lack a male reproductive tissue specific *LLG;* however, *A. comosus LLGG* is expressed at much higher levels than other *A. comosus LLG* paralogs in both male and female domains (Table S4). Hence, it is possible that *LLGG* in *A. comosus* is sufficient to perform the functions required in the male domain.

To obtain additional evidence in support of regulatory subfunctionalization of *LLG* expression in angiosperms, we examined *LLG* expression in Amborella, which is sister to all other extant angiosperms and contained only one *LLG.* In Amborella, the expression pattern of the sole *LLG* present (Figure 1) encompasses the expression domains of the other sampled *LLGs*. Interestingly, this single *LLG* showed vastly higher levels of expression in the male reproductive tissues compared to expression in the female or vegetative tissues (Table S4). Taken together, the expression analyses in 11 angiosperms showed that expansion of *LLG* gene family in angiosperms was accompanied by divergence in their expression and that regulatory subfunctionalization likely underlies the retention of duplicated *LLGs* in angiosperms.

### The inferred ancestral expression pattern of single copy angiosperm ancestor LLG is primarily in pollen and pollen tubes

Expression divergence among *LLGs* yielding copies that are primarily expressed in male reproductive tissues, while other gene copies are expressed in female reproductive tissues or more broadly raises the question: what was the ancestral expression pattern of *LLGs* in the earliest angiosperms? This question is also relevant given that the single copy *LLG* in Amborella showed vastly higher levels of expression in the male reproductive tissues. To better understand the evolution of divergent expression in *LLG* paralogs, we calculated normalized expression values (z-scores; Table S4) across the five expression domains for each *LLG* and used these values to reconstruct likelihood-based ancestral character states (Figures 6a and S4, Datasets S2 and S3). This expression reconstruction showed that a single copy *LLG* ancestor, with highest expression in male reproductive tissues, underwent independent expansions in monocots and eudicots.

**Figure 6.**
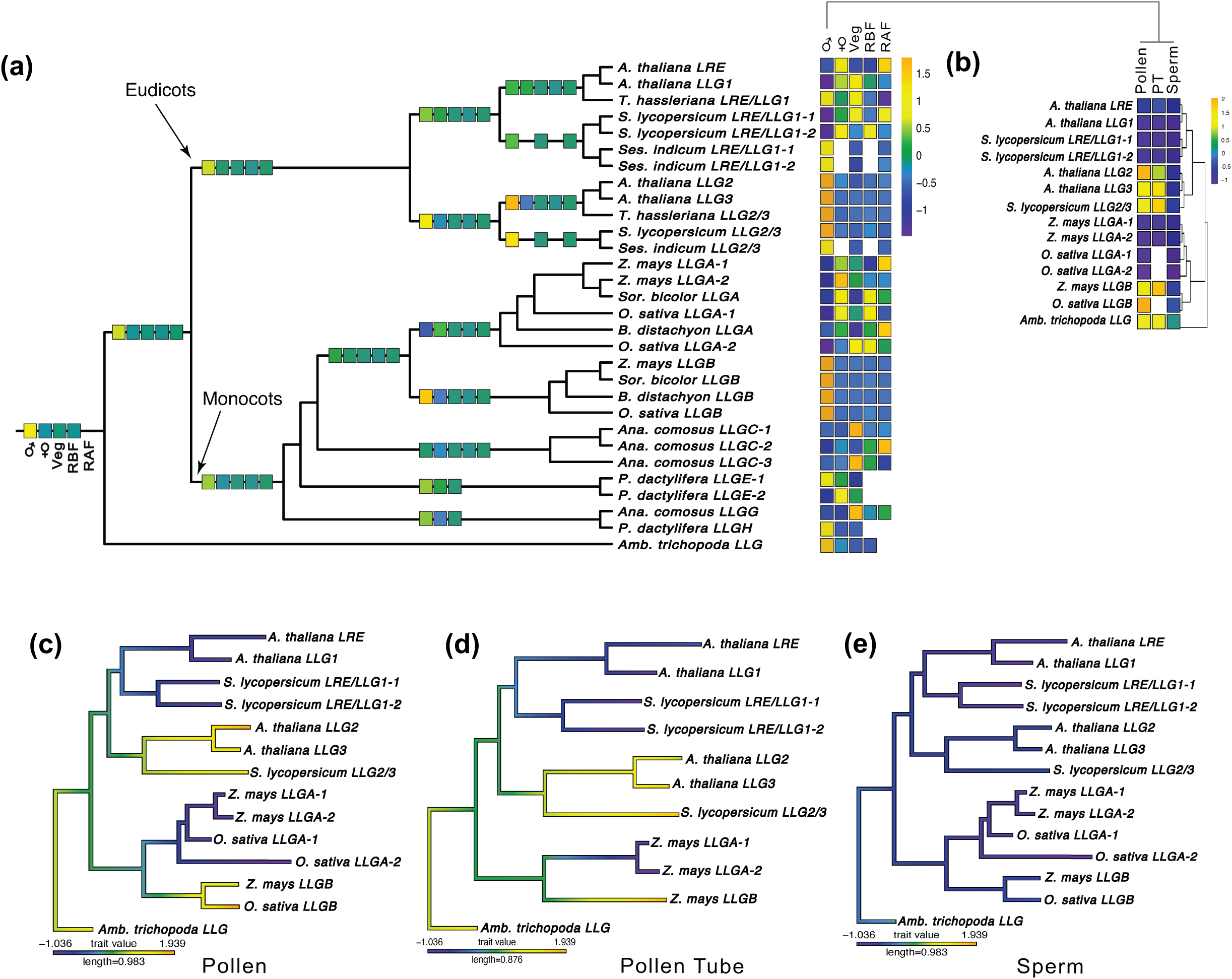
Phylogeny and expression analysis of *LLG* orthologs in angiosperms. (a) Maximum likelihood phylogenetic tree from putative ortholog CDS. The heatmap (right) displays the z-score expression of the orthologs in male, female, vegetative, reproductive tissues before fertilization (RBF), and reproductive tissues after fertilization (RAF). Ancestral state reconstruction was performed in phytools under either a Brownian motion or Ornstein-Uhlenbeck model. Reconstruction results of select nodes are displayed on the adjacent branches of the tree. Nodes with missing expression values are the result of missing expression data at the tips, precluding robust inference of the ancestral state at that node. (b) Heatmap of core male tissues for *LLGs* from a subset of species. The tissues shown are mature pollen, pollen tube (PT), and sperm respectively. (c-e) Ancestral state reconstruction trees for pollen, pollen tube, and sperm respectively. Tip expression and node expression are colored according to the z-score scales presented.

Male reproductive tissue comprises pollen grain, pollen tube, and sperm cells. To pinpoint relative expression level of *LLG*s within the male reproductive domain, we divided the male domain into pollen, pollen tubes, and sperm and then recalculated expression z-scores for each *LLG* present in five representative angiosperms from which expression data were available – two eudicots (Arabidopsis and *S. lycopersicum*), two monocots (*O. sativa* and *Z. mays*), and Amborella (Figure 6b, Dataset S4). These analyses revealed that expression is most pronounced in mature pollen and pollen tubes rather than sperm (Figure 6b, Dataset S4). Ancestral reconstruction of the expression in these tissues indicated that the single copy angiosperm *LLG* ancestor was likely highly expressed in pollen and pollen tube and expressed at low levels in sperm (Figures 6c–e, Datasets S2 and S3), pointing to critical functions for the ancestral *LLG*, and the signaling complex it was a part of, in male reproductive tissues.

## Discussion

### Regulatory Divergence likely led to the Retention of LLGs in Angiosperms

Multiple mechanisms have been proposed to explain the retention of duplicated genes, as progressive degeneration of one member of a paralogous set of genes is the default outcome (non-functionalization; (Panchy et al., 2016). Duplicated genes may acquire new functions (neo-functionalization) and hence may be retained (Ohno, 1970). Another mechanism of retention is sub-functionalization via sequence variations in the protein-coding or regulatory regions of the genes, by which the ancestral functions could be shared between the duplicated copies, resulting in non-overlapping expression of each paralog (Force et al., 1999). The advent of high-throughput sequencing and large-scale transcriptomic studies have allowed evaluation of expression divergence in duplicated genes in *A. thaliana* (Duarte et al., 2006; Zou et al., 2009; Liu et al., 2011; De Smet et al., 2017), *Glycine max* (Roulin et al., 2013), and *Zea mays* (Hughes et al., 2014). These studies used expression analysis to generate critical evidence in support of regulatory sub-functionalization and/or regulatory neo-functionalization. Here, we performed the first phylogenetic analysis of *LLGs* in angiosperms coupled with expression and functional studies of *LLG* gene family to show that regulatory subfunctionalization is likely the basis of retention of *LLG* duplicates.

In many organisms, there is a positive correlation between expression divergence and *cis*-regulatory motif divergence among duplicate genes (Li et al., 2005), including Arabidopsis (Arsovski et al., 2015). Consistent with this, we identified considerable variation among putative transcription factor binding sites in the putative promoter regions of *LLGs* (Figure S5). Additionally, in support of their shared expression in male reproductive tissues, specific transcription factor binding sites are conserved in the promoter regions of *LLG2* or *LLG3* of Arabidopsis and the single copy ortholog *LLG2/LLG3* in *C. violacea* (Figure S5). Similarly, a different set of transcription factor binding sites are conserved in the promoter regions of *LLG1* or *LRE* of Arabidopsis and the single copy ortholog *LRE/LLG1* in *C. violacea*, which show expression in female reproductive and vegetative tissues (Figure S5). Therefore, a conserved set of *cis*-regulatory elements is likely responsible for these divergent expression patterns in both Brassicaceae and Cleomaceae and ultimately contributed to the retention of duplicates in eudicots.

Preferential expression of at least one *LLG* copy in male reproductive tissues and highest expression of the single copy *LLG* in Amborella in the male domain raises the possibility of transcription factor-binding motifs that are conserved among the putative promoter regions of these genes. Using a set of *in vitro* binding data (see Methods), there was a clear pattern of presence of motifs in the *LLG2/3* clade in eudicots, as they shared AP2 (APETALA2) and EREBP (ethylene-responsive element binding proteins) transcription factor binding sites (Chandler and Werr, 2020) with the single copy Amborella *LLG* (Dataset S5). These differences are consistent with the distinct evolutionary history and divergence between the *LLGs* in eudicots and monocots. It is likely that the monocots have lost these motifs since their divergence from eudicots. In addition, it is likely that Arabidopsis *LLG2* and *Sesame indicum LRE/LLG1-2* have also lost these motifs compared to other orthologs (Dataset S5). Still, there is not one motif that is completely conserved among these genes (Dataset S5) and perhaps uncharacterized novel motifs are shared by these male tissue-expressed genes.

### The LLG Gene Family may be Co-evolving with the CrRLK1L and RALF Gene Families

Like *LLGs,* genes encoding the other two members of the trimeric complex (*Cr*RLK1Ls and RALFs), also show expression divergence in Arabidopsis (Campbell and Turner, 2017; Ge et al., 2017) and given our results, it is possible that they show similar divisions of expression domains in other angiosperms. The large size of *RALF* and *CrRLK1L* gene families pose a challenge in efforts to study all the possible combinations of the signaling complex (Kessler et al., 2015; Sharma et al., 2016). However, phylogenetic analysis coupled with functional and expression studies, like that performed in this study, may offer a viable approach to understand the evolution of expression domains in *CrRLK1L* and *RALF* families in angiosperms.

FER functions with both LRE and LLG1, and correspondingly, *FER* is expressed in both *LRE* and *LLG1* expression domains (Tsukamoto et al., 2010; Li et al., 2015). Given that specific *Cr*RLK1Ls, RALFs, and LLGs play distinct biological roles, *CrRLK1Ls* and *RALFs* may have co-evolved with members of the *LLG* family to perform their functions in different tissues. *In vitro* interactions of RALF23 with LLG1, LLG2, and LLG3 but not LRE provide support for this possibility (Xiao et al., 2019). Methods such as evolutionary rate covariation have been used to link co-evolution with functional associations (Findlay et al., 2014). Such methods, in combination with the phylogenetics, expression analyses, and molecular genetic assays used in this study will prove invaluable in further characterizing members of this critical signaling complex.

### The inferred ancestral Expression Pattern in Angiosperms Suggests an Important Role for LLG in the Evolution of Rapid Pollen Tube Growth During Angiosperm Reproduction

Results of our ancestral *LLG* expression reconstruction suggest that a signaling complex containing the single copy *LLG* may have mediated pollen tube growth in the earliest angiosperms. Moreover, the evolution of a faster reproductive cycle is often posited as a key innovation of early angiosperms (Stebbins, 1974). Several traits are believed to influence this increase in reproductive rate, including reduced male and female gametophytes and closed carpels. Moreover, evolution of faster pollen tube growth has been identified as a key link between the reduction in gametophyte size and closed carpels (Williams, 2008; Williams, 2012). Faster pollen tube growth in angiosperms is hypothesized to be facilitated by the accumulation of esterified pectins in pollen tube tips and deesterified pectins and callose in their walls (Williams, 2008). Consistent with this hypothesis, Arabidopsis *llg2 and llg3* mutant pollen tubes showed decreased length *in vivo* and *in vitro,* increased accumulation of esterified and deesterified pectins, and decreased levels of callose (Feng et al., 2019). Interestingly, RNA-seq-based transcriptome analyses of early land plants (Julca et al., 2021) revealed male-preferential expression of one of the three *LLG* homologs from the moss *Physcomitrella patens*, while maximal expression of both *LLG* homologs in the liverwort *Marchantia polymorpha* is in antheridia (Figure S6). Thus, it is possible that proto angiosperms co-opted an existing male expressed molecular signaling module, present across land plants, to regulate pollen tube growth through the carpel, thereby allowing relatively rapid delivery of sperm to the female gametophyte and setting the stage for the evolution of the suite of traits that unite all angiosperms.

## Materials and Methods

### Homolog Identification

Putative orthologs and paralogs from species representing major lineages of angiosperms were retrieved by first identifying conserved motifs in the four coding sequences (CDS) of *LLG* family members in the Arabidopsis genome. Homolgos were searched using CDS obtained from Phytozome v. 13.0 (Goodstein et al., 2012) or from the Database Resources of the National Center for Biotechnology Information (NCBI) (https://www.ncbi.nlm.nih.gov/) in MEME Suite v. 5.1.1 (Bailey, 1994) (Dataset S1). CDS of putative orthologs for *Cleome violacea* and *Tarenaya hassleriana* were obtained through the Comparative Genomics (CoGe) Platform using CoGe BLAST (tBLASTx) with *A. thaliana LRE* or *LLG1* nucleotide CDS as query (CoGe version (Altschul et al., 1990; Lyons and Freeling, 2008; Lyons et al., 2008)). Motif results and CDS were input into MAST (Bailey and Gribskov, 1998) under default parameters and only those with an E-value ≤ 1e-05 were retained as putative *LLG*s. Putative *LLGs* were edited to the longest possible ORF.

### Alignment and Phylogenetic Analyses

Based on preliminary alignments made using MUSCLE v. 3.8.425 (Edgar, 2004) in Geneious 2019.2.1 (https://www.geneious.com) under default parameters, we inferred maximum likelihood trees with RAxML-HPC MPI AVX2 v. 8.2.11 (Stamatakis, 2014). We used these preliminary trees to further assess the homology of sequences returned via motif searching and eliminated sequences whose homology could not be confidently assigned due to phylogenetic position, branch length, and/or increase to the length of the alignment.

The full amino acid sequences of putative *LLG* family members were aligned in MUSCLE v. 3.8.425 (Edgar, 2004). Using the Arabidopsis *LLG* gene structures as guides (Liu et al., 2016), the initial alignment was divided into signal peptide, main chain, and GPI anchor region. The alignment for each region was corrected by eye before being concatenated into a full-length amino acid alignment and submitted to TranslatorX (Abascal et al., 2010) along with the FASTA nucleotide sequences to obtain an amino acid corrected nucleotide alignment. The full alignment was partitioned and subjected to maximum likelihood phylogenetic analyses in RAxML-HPC MPI AVX2 v. 8.2.11 (Stamatakis, 2014) under the GTR+GAMMA model of sequence evolution. Support for branches of the tree was assessed from 1000 bootstrap replicates of the rapid bootstrapping algorithm in RaxML.

### Plant Growth

Arabidopsis (Columbia ecotype, Col-0) and *C. violacea* seeds were sterilized with 50% bleach and plated on 1/2X Murashigi and Skoog (MS) plates (Carolina Biological Supply Co.), with 1% sucrose for seedling growth assays and with corresponding antibiotics for transmission assays. After stratification (3 days in dark and 4 °C), plated seeds were incubated at 21 °C with continuous light (75–100 µmol·m^-2^·s^-1^). Ten-to fourteen-day-old seedlings were transplanted to soil and grown in 16 h light (100–120 µmol·m^-2^·s^-1^) at 21°C and 8 h dark at 18 °C.

### Cloning Transgenic Constructs

The *pLLG1::LRE-HA-GPI* and *pLLG1::LLG3* transgenic constructs were prepared using the primers and DNA templates (Table S4), and with the Arabidopsis *LLG1* promoter region (∼2 kb upstream of ATG) (Li et al., 2015). For SP-mRFP-LLG1 expression *in planta*, the Arabidopsis *LLG1* promoter was amplified from seedling genomic DNA using primers “ccaagcttgcatgccGTCGTTGTCCCAGATTCGTCG” and “gatctagagtcgaccGGTTCTTTGTTGGTTACAGGAGAAGTCAC” for subsequent cloning into the Gateway destination vector pGWB1 (Nakagawa *et al*., 2007a) using Infusion cloning (Takara Bio) and the SdaI/SbfI restriction enzyme (Thermo Scientific). The *SP-mRFP-LLG1* construct was synthesized with attached attB1/attB2 sites (Twist Bioscience) and subsequently recombined into pDONR-Zeo using BP II Clonase (Invitrogen). The resulting pDONR-Zeo SP-mRFP-LLG1 construct was recombined with *pGWB1-pLLG1*. The *pAtLLG1::SP-mRFP-LLG2* construct was synthesized (Thermo Scientific) with attB1/attB2 sites added and cloned into *pDONR-Zeo* via BP II Clonase (Invitrogen) and was subsequently recombined with *pGWB601* (Nakagawa et al., 2007) for *in planta* expression.

The *pAtLRE::LLG1-cYFP, pAtLRE::GUS*, *pAtLRE::Clevi-LRE/LLG1-cYFP,* and *pAtLLG1::Clevi-LRE/LLG1-cYFP* transgenic lines were created by replacing the genomic sequence of *LRE* in the *pAtLRE::LRE-cYFP* construct with the desired gene and/or promoter (Liu et al., 2016). PCR amplified products (PrimeSTAR GXL DNA Polymerase, TaKaRa Bio Inc.) using primers and DNA templates listed in Table S4 were cloned into *pLRE::LRE-cYFP* plasmid linearized with *SpeI-HF* and *AscI* by using the In-Fusion HD Cloning Plus system (Clontech). Transformants in Col-0 background (*pLRE::GUS*) or *lre-7* mutant (*pLRE::LRE-cYFP*, *pLRE::LLG1-cYFP*, and *pLRE::Clevi-LRE/LLG1-cYFP*) were generated and single-locus insertion lines were isolated (Liu et al., 2016).

The *pLLG1::SP-mRFP-LLG1* and *pLLG1::SP-mRFP-LLG2* lines were transformed into *llg1-2* plants by floral dip (Clough and Bent, 1998), and transformants were selected on 1x MS-agar with 1% sucrose, supplemented with 25 µg/mL kanamycin or 10 µg/mL glufosinate ammonium, respectively. Seedlings for seedling growth inhibition assays were grown in 12 h light (120 120 µmol·m^-2^·s^-1^) at 19–21 °C. Plants for ROS burst assays were grown in 10 h light (150 µmol·m^-2^·s^-1^) at 20 °C.

### cYFP and Seed Set Scoring

cYFP expression in mature unpollinated pistils were scored and imaged (Liu *et al*., 2016). Unfertilized, viable (enlarged after fertilization), and aborted ovules in siliques (10 days after pollination) were scored (Noble and Palanivelu, 2020). Three to five self-pollinated siliques located between 5th and 15th siliques from the bottom of the main branch of an Arabidopsis plant were scored.

### Root Hair and Hypocotyl Length Phenotypes in llg1 Seedlings

Root hair analysis was performed (Duan et al., 2010). Root hairs located between 1.5 and 3.5 mm from the primary root tip of four-day-old seedlings were observed with a stereoscope. The number of normal and defective (stunted and collapsed) root hairs was scored. The hypocotyl length assay was performed (Li et al., 2015). Three-day-old dark-grown seedlings were imaged using Epson Perfection V370 Photo Scanner at 600 dpi resolution and the length of hypocotyl was measured with Image J.

### Propidium Iodide Staining of Pavement Cells

Six-day-old seedlings were stained with 50 μg/mL propidium iodide for 20 min and washed with ddH_2_O. Images were acquired using confocal microscopy on a NIKON A1 Spectral System (excitation wavelength 561 nm and emission wavelength 595 nm) and analyzed by the NIS-Elements AR Analysis Software (V 5.02).

### RALF1-induced Root Growth Inhibition Assays

The root sensitivity assays with RALF1 treatment were performed (Haruta et al., 2014; Li et al., 2015): three-day-old light-grown seedlings were treated with RALF1 for 2 days at concentrations indicated in Figures 3 and S2. Growth during treatment was determined by measuring primary root length using Image J at the beginning and end of treatments.

### RALF23-induced Seedling Growth Inhibition Assays

Five-day-old T2 *pLLG1::SP-mRFP-LLG1* and *pLLG1::SP-mRFP-LLG2* T2 seedlings were transferred to liquid 1x MS with 1% sucrose in sterile 48-well plates, with or without 1 µM RALF23 peptide (Scilight) (Xiao et al., 2019). Seedling fresh weight was measured after 7 days of growth in liquid medium.

### ROS Burst Measurements

ROS burst measurements were performed (Xiao et al., 2019). Twelve 4-mm leaf discs from 4.5-week-old plants were harvested and equilibrated overnight in 96-well plates in sterile, deionized water. The next day, water was replaced with PAMP solution (100 nM flg22 or elf18 peptide plus 10 µg/mL horseradish peroxidase and 100 µM luminol). ROS was immediately measured using a charge-coupled device camera (Photek). Total ROS production was calculated as the sum of Random Light Units value over 40 minutes of PAMP treatment.

### GUS Staining

Tissues were stained for GUS activity (Johnson and Kost, 2010), mounted in 50% glycerol, observed on a Zeiss Axiovert 100 epifluorescence microscope, and imaged using Metamorph (Version 7; Molecular Devices). For each tissue and genotype, at least five samples were observed.

### Transcription Factor Binding Site Analysis

DAP-seq data were obtained from the Plant Cistrome Database (http://neomorph.salk.edu/dev/pages/shhuang/dap_web/pages/index.php) in the MEME motif file format. These 838 motifs were scanned against the putative promoter sequences (1 kb upstream of the transcriptional start, not overlapping with neighboring genes) of the *LLG* paralogs with MAST (Bailey and Gribskov, 1998) using default parameters. The scanned results were used to determine the presence of motif sites (mapping p-value was <1e-3). The motif site information was combined for all motifs to generate a motif presence (1)/absence (0) matrix for hierarchical clustering using the heatmap.2 function in R.

### Expression Analyses

TPM (transcript per kilobase million) values from CoNekT (https://evorepro.sbs.ntu.edu.sg/) for Arabidopsis, *S. lycopersicum*, *Z. mays*, *O. sativa*, and Amborella (Mutwil et al., 2011; Ruprecht et al., 2017; Julca et al., 2021). For *LLGs* (Dataset S1) in *S. bicolor* and *B. distachyon* (Davidson *et al*., 2012), TPM values were obtained from the EMBL-EBI expression database (https://www.ebi.ac.uk/gxa/home). FPKM (fragments per kilobase million) expression values reported for *T. hassleriana* (Kulahoglu et al., 2014) and *A. comosus* (Wang et al., 2020) were converted to TPM by dividing the FPKM value for each gene by the sum of FPKM values for all genes, then multiplying that number by one million. For *S. indicum* and *P. dactylifera* raw RNA-seq data were retrieved from the NCBI Sequence Read Archive (SRA - https://www.ncbi.nlm.nih.gov/sra; Bioprojects PRJNA74261 and PRJNA472694, respectively). Both retrieval and mapping were performed in the CyVerse Discovery Environment (Merchant et al., 2016) using the RMTA pipeline v. 2.6.3 (Peri et al., 2019) under default parameters. Read count data for focal genes were then extracted and converted to TPM. We defined five expression domains based on available RNA-seq data: male, female, reproductive before and after fertilization, and vegetative tissues. We binned sampled tissues with TPM values into one of these five categories, calculated the mean TPM of each expression domain, and then z-transformed these values in R v. 4.0.2 (R Core Team, 2020). Expression within the male domain was obtained by calculating expression z-scores for pollen, pollen tube, and sperm for species in which these tissues were available in RNA-seq experiments.

### Ancestral Character State Reconstruction

We used the z-score of expression for each *LRE/LLG* sequence along with our phylogeny, pruned to only those family members from which expression data were available, to infer the ancestral level of relative expression at nodes in the tree. Ancestral state reconstruction was performed in R v. 4.0.2 (R Core Team, 2020). We used both Brownian Motion (BM) and Ornstein-Uhlenbeck (OU) models in phytools v0.7-47 (50) and then compared their log-likelihoods via Akaike information criterion (AIC) (Akaike, 1974) and AICc (Mazerolle, M. J. (2019), R package version 2.1-1. Retrieved from https://CRAN.R-project.org/package=AICcmodavg) (Dataset S2). The latter employs a correction and may be more accurate for data sets of the size we analyzed.

### Image Processing

ImageJ was used to assemble image panels and insert scale bars.

## Acknowledgements

We acknowledge Dr. Patrick Edger from Michigan State University for providing us access to the *Cleome violacea* genome on CoGe and Dr. Jocelyn Hall from the University of Alberta for providing *Cleome violacea* seeds. We thank Dr. Ramin Yadegari (University of Arizona) for the Zeiss Axiophot microscope. We thank Dr. Mark Johnson, Brown University and members of the Palanivelu Lab for insightful discussions. JAN was supported by the following: IGERT Comparative Genomics Program at the University of Arizona (Award ID: 0654435); NSF Graduate Research Fellowship: Grant DGE-1143953; the Boynton Graduate Fellowship; and the University of Arizona Graduate College Office of Diversity and Inclusion. NVB was supported by a University of Arizona Graduate College University Fellowship and an NSF grant to MAB (IOS-1546825). Additional support for this work was provided by an NSF grant to RP (IOS-1146090) and University of Arizona Undergraduate Biology Research Program fellowship to SH (private donors). This work was also supported by the Gatsby Charitable Foundation (C.Z.), the University of Zürich (C.Z.), the European Research Council under the European Union (EU)’s Horizon 2020 research and innovation programme (grant agreement No 773153, project ‘IMMUNO-PEPTALK’), as well as fellowships from the European Molecular Biology Organization, the Natural Sciences and Engineering Research Council of Canada and the Deutsche Forschungsgemeinschaft (fellowships EMBO-LTF 100-2017 and NSERC PDF-532561-2019 to T.A.D.; DFG STE 2448/1 to M.S.T.). Work in AYC lab was supported by Natural Science Foundation (IOS-1645854 and MCB-1715764) to AYC and Hen-Ming Wu, and the National Institute of Food and Agriculture, U.S. Department of Agriculture, the Center for Agriculture, Food, and the Environment under project number MAS00525. The contents are solely the responsibility of the authors and do not necessarily represent the official views of the USDA or NIFA. We also thank HMW for the contribution of LLG1 and LLG3 constructs. KM was supported by the Torrey Summer Research Scholarship from UMass Plant Biology Program and the Linda Slakey Summer Research Scholarship from UMass BMB department. We also acknowledge funding from NSF pollen RCN grant (MCB0955910) for sponsoring activities and meetings that forged collaborations between AYC and RP labs. Work in the ADLN lab was supported by the National Science Foundation (PGRP-IOS 2023310 and PGRP-IOS 1758532). Work in the Shiu lab was partly supported by the National Science Foundation (IOS-1546617, DEB-1655386) and U.S. Department of Energy (Great Lakes Bioenergy Research Center BER DE-SC0018409).

