## Supplementary Materials_Figures for "Evolutionary analysis of the LORELEI gene family in angiosperms reveals regulatory subfunctionalization"

### Supplemental Information

**Fig. S1. LRE-HA, LLG3, and Clevi-LRE/LLG1-cYFP complemented root hair lengths in *llg1-2* seedlings prior to RALF treatment and increased RALF1 sensitivity after treatment.**

Root lengths before and after RALF1 treatment were measured in single representative lines from *LRE-HA, LLG3,* and *Clevi-LRE/LLG1-cYFP*. Root lengths in three-day-old seedlings were measured, then treated with 0 μM RALF1 (untreated), 0.5 μM RALF1, or 1 μM RALF1. Roots were measured two days after RALF1 treatments with three trial replicates. Error bars represent ±SD.

**Fig. S2. ROS production kinetics in response to flg22 (top) or elf18 (bottom) is restored in *pLLG1::SP-mRFP-LLG1* or *pLLG1::SP-mRFP-LLG2* lines.**

ROS burst levels in response to 100 nM flg22 (top) or elf18 (bottom) over time is indicated in Relative Light Unit (RLU). In each trace of indicated genotypes, data shown is average of n=12 leaf discs, ±SE.

**Fig. S3. LLG1-cYFP and Clevi-LRE/LLG1-cYFP were expressed in synergid cells and localized to the FA.**

(A-B) Diagrams of the *pLRE::LLG1-cYFP* and *pLRE::Clevi-LRE/LLG1-cYFP* constructs. (C) A diagram of a mature ovule with a 7-celled female gametophyte. Synergid cells are located in the micropylar end of the ovule, adjacent to the egg cell. The finger-like projections of the FA are shown in yellow. A red arrow points to the FA. (D) In mature unpollinated pistils, LRE-cYFP is expressed in the synergid cells, with localization in the puncta in the synergid cell cytoplasm and in the FA. The ovule is outlined in light gray dashed line, while the female gametophyte is outlined in dark gray dashed line. The red rectangle marks the synergid cells. (E) Close-up image of the LRE-cYFP in the synergid cells marked by the red rectangle in Fig C, outlined in dark gray dashed line. YFP localized in the FA and puncta in the synergid cells. (F) Close-up image of the LLG1-cYFP in the synergid cells with YFP localization in the puncta and the FA of the synergid cells. (G) Close-up image of the Clevi-LRE/LLG1-cYFP in the synergid cells. YFP is weakly expressed in the FA, but is not present elsewhere in the synergid cells, including the puncta.

**Fig. S4. LRE/LLG phylogeny with labeled nodes from ancestral state reconstruction analysis.**

Nodes from ancestral state reconstruction analysis are numbered 1-29. Node labels can be compared with Dataset S4 to retrieve reconstructed ancestral expression z-scores for each tissue. Nodes discussed in the main text are named.

**Fig. S5. Distribution of putative transcription factor (TF) binding sites in the putative promoter regions of LRE homologs in *Arabidopsis thaliana* and *Cleome violacea*.**

(A) TFs with putative binding sites as determined using DNA affinity purification-sequencing (DAP-seq) are shown as present (yellow boxes, $p < 1e\text{-}4$) or absent (blue boxes). The TF family and gene name information were based on the Plant Cistrome Database. *: amplified DAP-seq where secondary DNA modifications were removed. (B) Table showing the frequency of putative TF binding site occurrence in the promoters of *LRE*, *LLG1*, and the single copy ortholog *LRE/LLG1* in *Cleome violacea.* (C) Table showing the frequency of putative TF binding site occurrence in the promoters of *LLG2*, *LLG3*, and the single copy ortholog *LLG2/LLG3* in *Cleome violacea.*

**Fig. S6. A comparative heat map of *LLG* expression in different tissues of *Marchantia polymorpha* and *Physcomitrella patens*.**

RNA-seq generated raw normalized (expression values normalized per gene) expression values of *LLG* genes in the indicated tissues of *Marchantia polymorpha* and *Physcomitrella patens* were compared to generate the heat map in the CoNekT database. Red and green cells indicate high and low expression, respectively. Lack of expression values are indicated by dark gray cells.

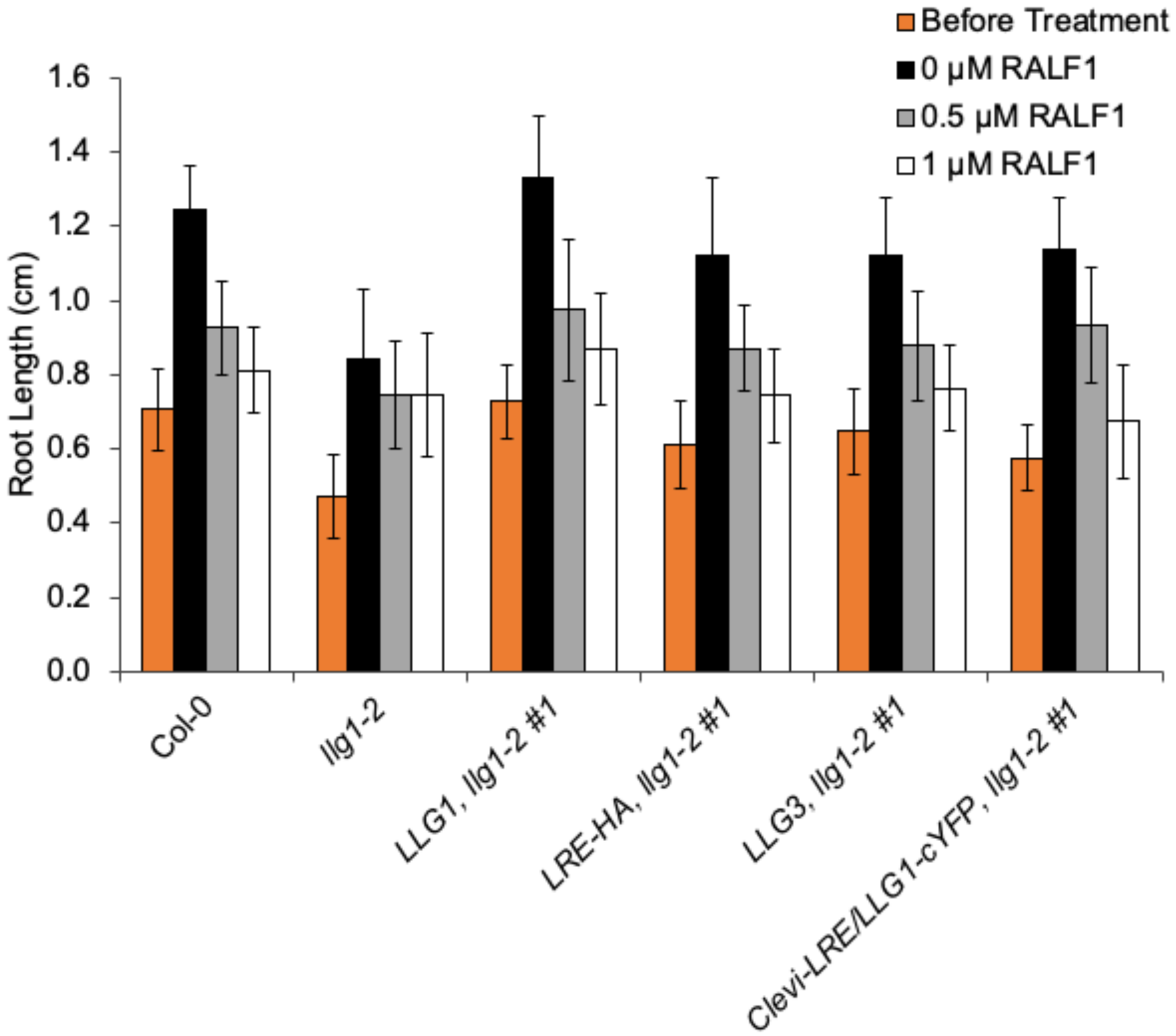

Before Treatment
0 μM RALF1
0.5 μM RALF1
1 μM RALF1
Root Length (cm)
1.6
1.4
1.2
1.0
0.8
0.6
0.4
0.2
0.0
Col-0
llg1-2
LLG1, llg1-2 #1
LRE-HA, llg1-2 #1
LLG3, llg1-2 #1
Clevi-LRE/LLG1-cYFP, llg1-2 #1


**Fig. S1**

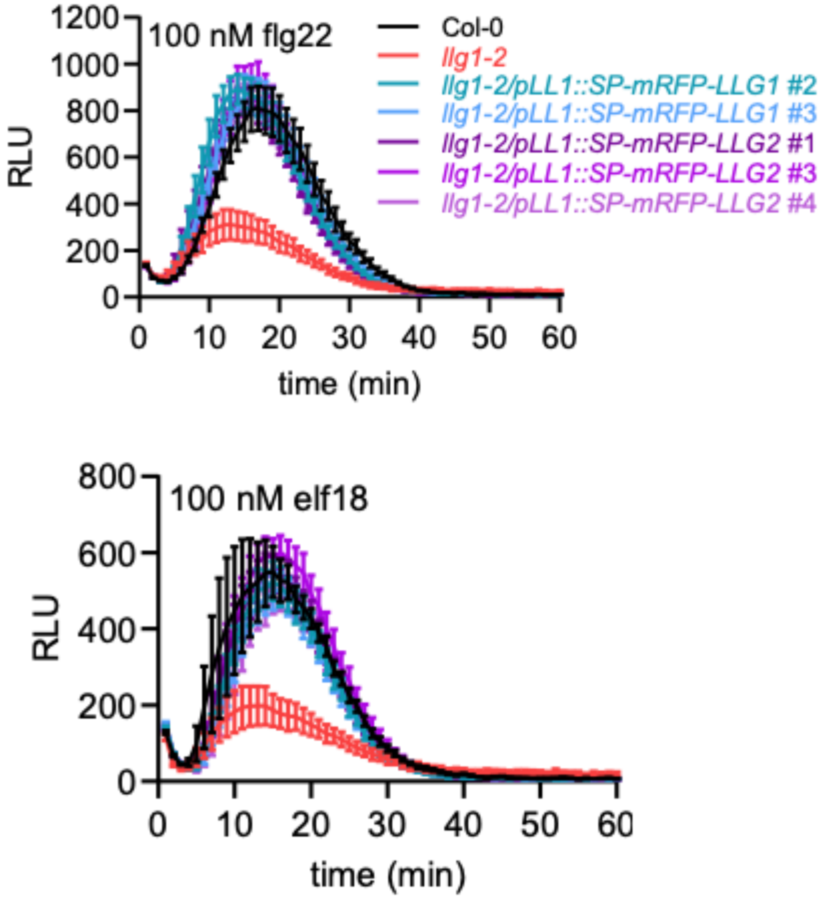
100 nM flg22
Col-0
llg1-2
llg1-2/pLL1::SP-mRFP-LLG1 #2
llg1-2/pLL1::SP-mRFP-LLG1 #3
llg1-2/pLL1::SP-mRFP-LLG2 #1
llg1-2/pLL1::SP-mRFP-LLG2 #3
llg1-2/pLL1::SP-mRFP-LLG2 #4
RLU
time (min)
100 nM elf18
RLU
time (min)

**Fig. S2**

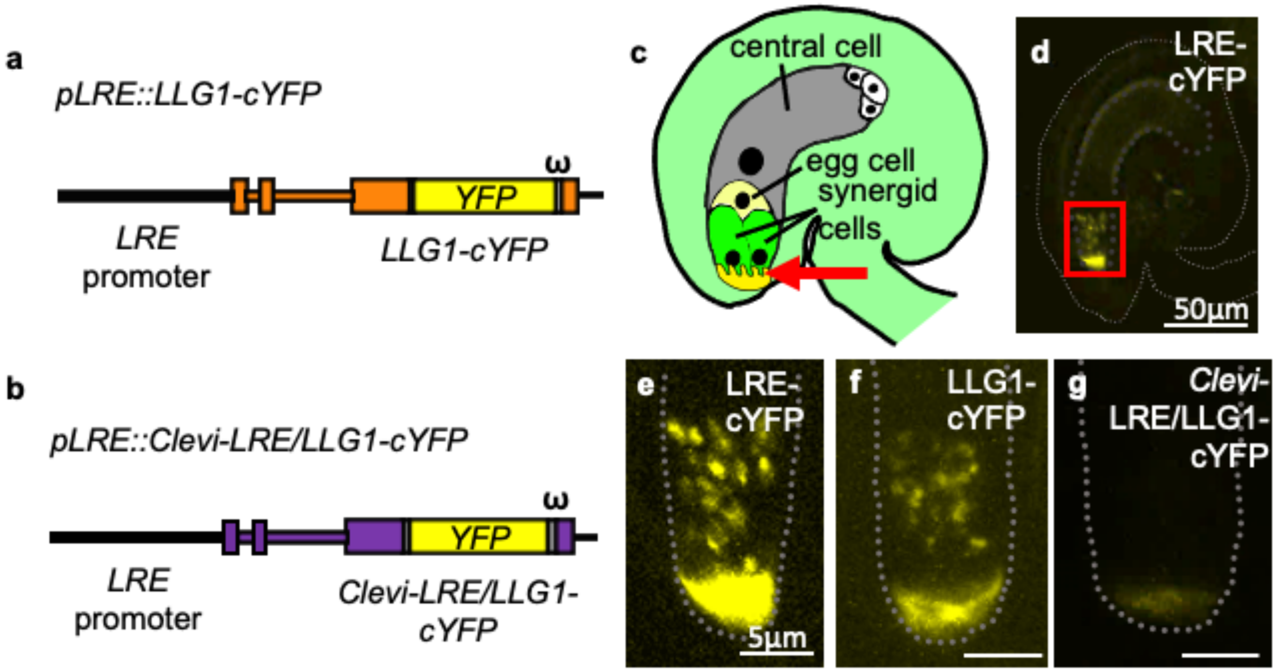
a
pLRE::LLG1-cYFP
ω
YFP
LRE
promoter
LLG1-cYFP
b
pLRE::Clevi-LRE/LLG1-cYFP
ω
YFP
LRE
promoter
Clevi-LRE/LLG1-
cYFP
c
central cell
egg cell
synergid
cells
d
LRE-
cYFP
50µm
e
LRE-
cYFP
5µm
f
LLG1-
cYFP
g
Clevi-
LRE/LLG1-
cYFP

Fig. S3

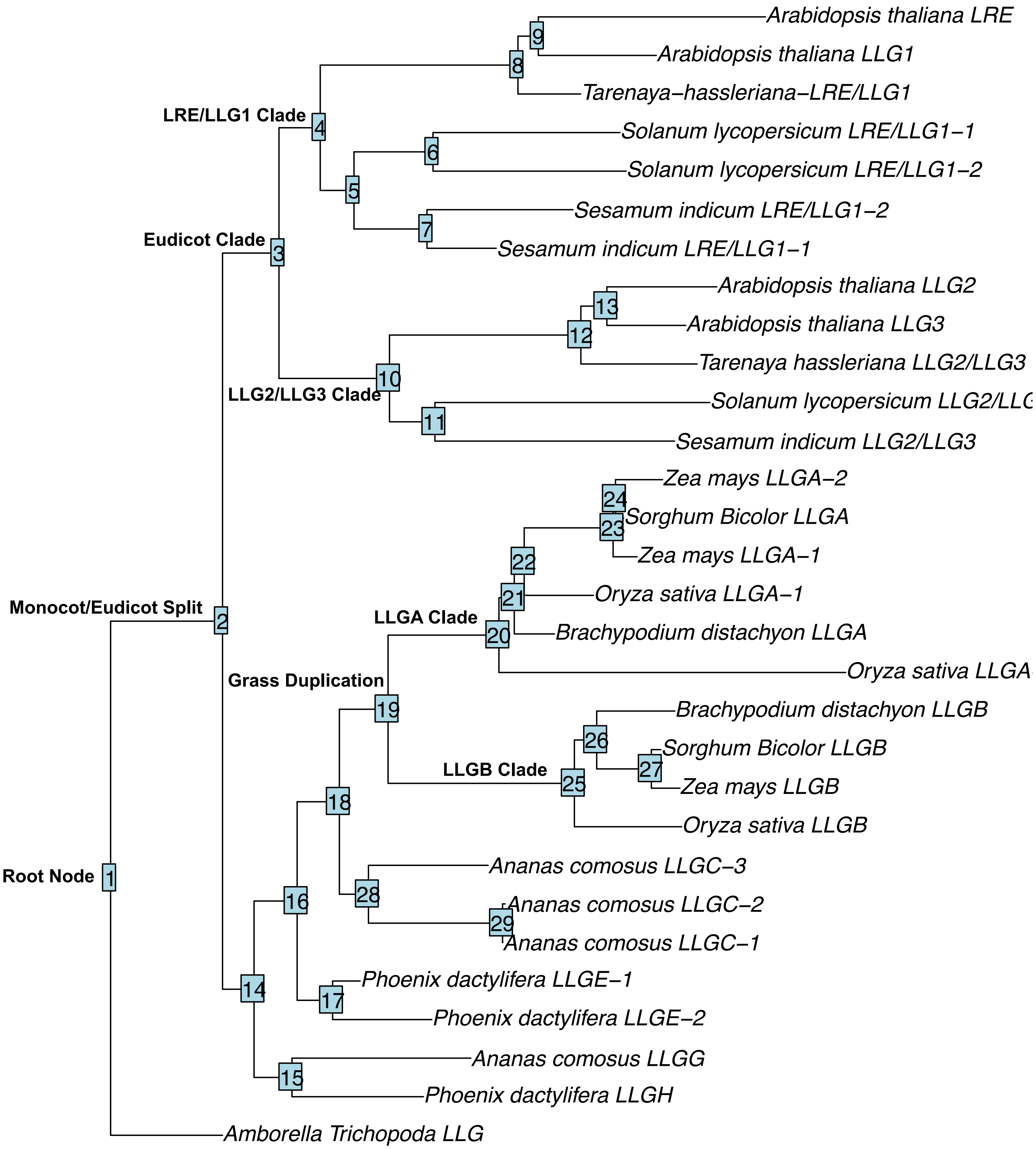
Arabidopsis thaliana LRE
Arabidopsis thaliana LLG1
Tarenaya−hassleriana−LRE/LLG1
LRE/LLG1 Clade
Solanum lycopersicum LRE/LLG1−1
Solanum lycopersicum LRE/LLG1−2
Sesamum indicum LRE/LLG1−2
Eudicot Clade
Sesamum indicum LRE/LLG1−1
Arabidopsis thaliana LLG2
Arabidopsis thaliana LLG3
Tarenaya hassleriana LLG2/LLG3
LLG2/LLG3 Clade
Solanum lycopersicum LLG2/LL
Sesamum indicum LLG2/LLG3
Zea mays LLGA−2
Sorghum Bicolor LLGA
Zea mays LLGA−1
Oryza sativa LLGA−1
Monocot/Eudicot Split
LLGA Clade
Brachypodium distachyon LLGA
Oryza sativa LLGA
Grass Duplication
Brachypodium distachyon LLGB
Sorghum Bicolor LLGB
LLGB Clade
Zea mays LLGB
Oryza sativa LLGB
Ananas comosus LLGC−3
Root Node
Ananas comosus LLGC−2
Ananas comosus LLGC−1
Phoenix dactylifera LLGE−1
Phoenix dactylifera LLGE−2
Ananas comosus LLGG
Phoenix dactylifera LLGH
Amborella Trichopoda LLG

(a)

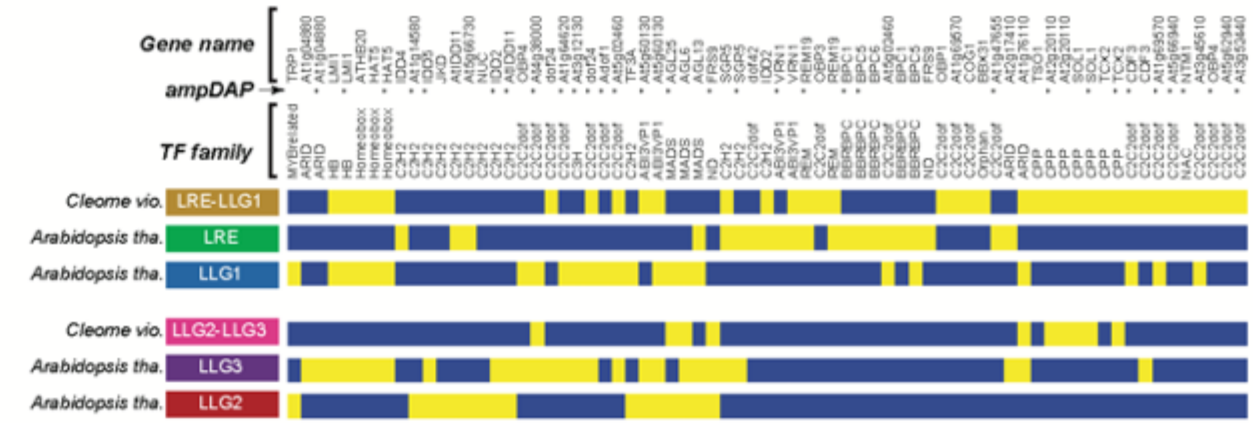


(b)

| Motif present in promoters of | Type occurrence | % occurrence |
|---|---|---|
| None | 18 | - |
| *LLG1* only | 9 | 14.52 |
| *LRE* only | 13 | 20.97 |
| *Clevi LRE/LLG1* only | 20 | 30.65 |
| *LLG1* & *LRE* | 3 | 4.84 |
| *LLG1* & *Clevi LRE/LLG1* | 12 | 19.35 |
| *LRE* & *CleviLRE/LLG1* | 5 | 8.06 |
| All three | 0 | 0.00 |
| Total | 80 | - |
| Once in any of the 3 | 62 | - |

(c)

| Motif present in promoters of | Type occurrence | % occurrence |
|---|---|---|
| None | 40 | - |
| *LLG2* only | 17 | 42.5 |
| *LLG3* only | 7 | 17.5 |
| *Clevi LRE/LLG1* only | 5 | 12.5 |
| *LLG2* & *LLG3* | 6 | 15 |
| LLG2 & *Clevi LRE/LLG1* | 2 | 5 |
| *LLG3* & *Clevi LRE/LLG1* | 1 | 2.5 |
| All three | 2 | 5 |
| Total | 80 | - |
| Once in any of the 3 | 40 | - |

**Fig. S5**

| | Gene | Egg cell/ovule/archegonia | Pollen/sperm/antheridia | Flowers | Seedlings | Meristem | Seeds/embryo/spores | Stems | Leaves | Roots |
|---|---|---|---|---|---|---|---|---|---|---|
| ***Marchantia polymorpha*** | Mapoly0048s0110 | 0.79 | 1.0 | | | | 0.52 | | | |
| | Mapoly0090s0020 | 0.53 | 1.0 | | | | 0.85 | | | |
| ***Physcomitrella patens*** | Pp3c4_4130V1.1 | 0.74 | 0.77 | | | | 0.31 | | 1.0 | |
| | Pp3c4_4140V1.1 | 0.0 | 0.0 | | | | 1.0 | | 0.0 | |
| | Pp3c8_1150V1.1 | 0.0 | 0.07 | | | | 1.0 | | 0.73 | |

**Fig. S6**
