## Supplementary Materials_Tables for "Evolutionary analysis of the LORELEI gene family in angiosperms reveals regulatory subfunctionalization"

**Table S1.** Enhanced transmission of the *pLRE::LLG1-cYFP* transgene through the *lre-7* female gametophyte.

| Parents | | Observed No. of progeny | | Transmission Efficiency (TE) Analysis | | |
|---|---|---|---|---|---|---|
| Female parent+ | Male parent+ | Hyg$^{R*}$ | Hyg$^{S*}$ | TE (R/S) | χ2† | P-value |
| *WT* | *LRE-cYFP-23* | 129 | 116 | 1.11 | 0.34 | 0.56# |
| *LRE-cYFP-23* | *WT* | 156 | 31 | 5.03 | 47.13 | 6.64E-12 |
| *WT* | *LLG1-cYFP-A2* | 213 | 171 | 1.25 | 2.30 | 0.13# |
| *LLG1-cYFP-A2* | *WT* | 163 | 24 | 6.79 | 60.05 | 9.25E-15 |
| *WT* | *LLG1-cYFP-10* | 60 | 71 | 0.85 | 0.46 | 0.50# |
| *LLG1-cYFP-10* | *WT* | 179 | 30 | 5.97 | 60.95 | 5.87E-15 |
| *WT* | *LLG1-cYFP-11* | 147 | 79 | 1.86 | 10.47 | 0.001# |
| *LLG1-cYFP-11* | *WT* | 138 | 16 | 8.63 | 57.32 | 3.71E-14 |

+ Line Numbers refer to three independent transformants in the *lre-7/lre-7* background containing single insertion of the *pLRE::LLG1-cYFP* transgene. Genotype of each transgenic line is heterozygous for the transgene (*pLRE::LLG1-cYFP/+*) and homozygous for the *lre-7* mutation (*lre-7/lre-7*).

***** Hygromycin resistant (Hyg$^{R}$) and susceptible (Hyg$^{S}$) progeny. Hygromycin resistance gene is linked to the construct carrying the *pLRE::LLG1-cYFP* transgene.

TE, Transmission efficiency was calculated as the ratio of hygromycin resistance (R) to susceptibility (S) in the progeny of the indicated cross

† χ2 is calculated based on an expected segregation ratio of hygromycin resistant to susceptibility of 1:1

### No significant deviation from 1:1 segregation through the male gametophyte indicates that pollen parent contains a single insertion of the *pLRE::LLG1-cYFP* transgene. Additional details on our protocol to isolate single insertion lines can be found in the methods.

**Table S2.** Enhanced transmission of the *pLRE::Clevi-LRE/LLG1- cYFP* transgene through the *lre-7* female gametophyte.

| Parents | | Observed No. of progeny | | Transmission Efficiency (TE) Analysis | | |
|---|---|---|---|---|---|---|
| Female parent+ | Male parent+ | $Hyg_R$* | $Hyg_S$* | TE (R/S) | χ2† | P-value |
| *WT* | *LRE-cYFP-23* | 212 | 182 | 1.16 | 1.14 | 0.28# |
| *LRE-cYFP-23* | *WT* | 335 | 43 | 7.79 | 132.5 6 | 1.13E-30 |
| *WT* | *Clevi-LRE/LLG1-cYFP-10* | 127 | 130 | 0.98 | 0.02 | 0.89# |
| *Clevi-LRE/LLG1-cYFP-10* | *WT* | 87 | 52 | 1.67 | 4.49 | 0.03 |
| *WT* | *Clevi-LRE/LLG1-cYFP-11* | 150 | 113 | 1.33 | 2.62 | 0.10# |
| *Clevi-LRE/LLG1-cYFP-11* | *WT* | 109 | 52 | 2.10 | 10.44 | 0.001 |
| *WT* | *Clevi-LRE/LLG1-cYFP-12* | 90 | 88 | 1.02 | 0.01 | 0.92# |
| *Clevi-LRE/LLG1-cYFP-12* | *WT* | 84 | 28 | 3 | 14.93 | 0.0001 |

\+ Line Numbers refer to three independent transformants in the *lre-7/lre-7* background containing single insertion of the *pLRE::Clevi-LRE/LLG1-cYFP* transgene. Genotype of each transgenic line is heterozygous for the transgene (*pLRE::Clevi-LRE/LLG1-cYFP/+*) and homozygous for the *lre-7* mutation (*lre-7/lre-7*).

***** Hygromycin resistant ($Hyg_R$) and susceptible ($Hyg_S$) progeny. Hygromycin resistance gene is linked to the construct carrying the *pLRE::Clevi-LRE/LLG1-cYFP* transgene.

TE, Transmission efficiency was calculated as the ratio of hygromycin resistance (R) to susceptibility (S) in the progeny of the indicated cross

† χ2 is calculated based on an expected segregation ratio of hygromycin resistant to susceptibility of 1:1

**Table S3.** Expression levels of single copy *LRE/LLG* ortholog and single copy *LLG2/LLG3* ortholog in Cleomaceae (as reported in Kuelahoglu et al. 2014)[a]

| Tissues | *Tarenaya hasseleriana* | | *Gynandropsis gynandra* | |
|---|---|---|---|---|
| | *LRE/LLG1* | *LLG2/LLG3* | *LRE/LLG1* | *LLG2/LLG3* |
| Seedling | 145 ± 51 | 0 ± 0 | 192 ± 44 | 2 ± 0 |
| Leaves | 117 ± 19 | 0 ± 0 | 90 ± 44 | 1 ± 1 |
| Roots | 176 ± 31 | 0 ± 0 | 98 ± 10 | 2 ± 0 |
| Stems | 173 ± 42 | 0 ± 0 | 120 ± 5 | 1 ± 0 |
| Sepals | 96 ± 19 | 1 ± 0 | 159 ± 12 | 2 ± 2 |
| Petals | 47 ± 10 | 1 ± 1 | 102 ± 12 | 3 ± 1 |
| Stamens | 164 ± 8 | 98 ± 36 | 77 ± 18 | 71 ± 9 |
| Carpels | 90 ± 2 | 0 ± 0 | 107 ± 28 | 0 ± 1 |
| Seedlings | 43 ± 22 | 0 ± 0 | 39 ± 13 | 3 ± 2 |

[a]The numbers in the table represent average RPKM (± Standard Deviation) of RNA seq reads obtained and normalized from 3 biological replicates of indicated tissues.

**Table S4.** Average expression of *LLGs* across five tissue domains

| *LLG* Gene names | Male | | Female | | RBF | | RAF | | Veg | |
|---|---|---|---|---|---|---|---|---|---|---|
| | Avg. TPM | z-score | Avg. TPM | z-score | Avg. TPM | z-score | Avg. TPM | z-score | Avg. TPM | z-score |
| Eudicots | | | | | | | | | | |
| *Arabidopsis thaliana LLG1* | 15.83 | -1.44 | 90.76 | 0.56 | 67.30 | -0.07 | 58.77 | -0.29 | 116.25 | 1.24 |
| *Arabidopsis thaliana LLG2* | 317.05 | 1.78 | 45.10 | -0.30 | 24.20 | -0.46 | 35.60 | -0.38 | 1.58 | -0.64 |
| *Arabidopsis thaliana LLG3* | 205.88 | 1.79 | 0.62 | -0.46 | 3.09 | -0.43 | 1.03 | -0.45 | 0.84 | -0.45 |
| *Arabidopsis thaliana LRE* | 0.13 | -0.61 | 0.99 | 0.70 | 0.02 | -0.79 | 1.46 | 1.41 | 0.07 | -0.71 |
| *Sesamum indicum LLG2/LLG3* | 270.33 | 1.15 | NA | NA | NA | NA | 13.16 | -0.53 | 0.00 | -0.62 |
| *Sesamum indicum LRE/LLG1-1* | 182.22 | 1.15 | NA | NA | NA | NA | 5.79 | -0.55 | 0.00 | -0.61 |
| *Sesamum indicum LRE/LLG1-2* | 24.97 | 1.13 | NA | NA | NA | NA | 5.88 | -0.34 | 0.00 | -0.79 |
| *Solanum lycopersicum LLG2/LLG3* | 164.54 | 1.78 | 8.86 | -0.41 | 17.33 | -0.29 | 0.09 | -0.54 | 0.09 | -0.54 |
| *Solanum lycopersicum LRE/LLG1-1* | 0.24 | -1.48 | 56.87 | 0.08 | 39.28 | -0.41 | 87.00 | 0.91 | 87.00 | 0.91 |
| *Solanum lycopersicum LRE/LLG1-2* | 0.50 | -1.37 | 54.24 | 1.06 | 50.93 | 0.91 | 24.18 | -0.30 | 24.18 | -0.30 |
| *Tarenaya hassleriana LLG2/LLG3* | 56.80 | 1.79 | 0.00 | -0.50 | 2.71 | -0.39 | 1.67 | -0.43 | 1.11 | -0.46 |
| *Tarenaya hassleriana LRE/LLG1* | 131.13 | 1.13 | 81.82 | -0.05 | 65.77 | -0.43 | 25.12 | -1.40 | 115.08 | 0.75 |
| Grass monocots (Poaceae) | | | | | | | | | | |
| *Brachypodium distachyon LLGA* | 44.00 | -0.54 | 75.00 | 0.20 | 70.00 | 0.08 | 128.25 | 1.48 | 16.00 | -1.21 |
| *Brachypodium distachyon LLGB* | 399.00 | 1.79 | 3.00 | -0.43 | 0.00 | -0.45 | 0.00 | -0.45 | 0.00 | -0.45 |
| *Oryza sativa LLGA-1* | 0.02 | -1.24 | 87.39 | 0.82 | 102.17 | 1.17 | 25.56 | -0.64 | 48.45 | -0.10 |
| *Oryza sativa LLGA-2* | 0.07 | -1.46 | 18.33 | -0.35 | 36.38 | 0.75 | 23.49 | -0.04 | 42.12 | 1.09 |
| *Oryza sativa LLGB* | 378.65 | 1.79 | 0.65 | -0.48 | 23.47 | -0.34 | 0.13 | -0.48 | 0.01 | -0.48 |
| *Sorghum Bicolor LLGA* | 36.00 | -0.86 | 115.00 | 1.17 | 97.00 | 0.70 | 76.00 | 0.16 | 24.00 | -1.17 |
| *Sorghum Bicolor LLGB* | 206.00 | 1.79 | 4.00 | -0.45 | 9.50 | -0.39 | 1.97 | -0.47 | 0.80 | -0.48 |
| *Zea mays LLGA-1* | 16.67 | -0.92 | 38.27 | 0.51 | 16.59 | -0.92 | 52.11 | 1.43 | 29.11 | -0.10 |
| *Zea mays LLGA-2* | 21.13 | -1.07 | 88.04 | 1.64 | 40.32 | -0.29 | 40.22 | -0.29 | 47.60 | 0.01 |
| *Zea mays LLGB* | 798.83 | 1.79 | 8.59 | -0.43 | 0.44 | -0.46 | 0.06 | -0.46 | 6.83 | -0.44 |
| Non grass monocots | | | | | | | | | | |
| *Ananas comosus LLGC-1* | 1.92 | -0.48 | 1.22 | -0.62 | 2.47 | -0.37 | 2.84 | -0.30 | 13.52 | 1.78 |
| *Ananas comosus LLGC-2* | 11.29 | -0.89 | 15.47 | -0.25 | 18.32 | 0.18 | 27.81 | 1.63 | 12.69 | -0.68 |
| *Ananas comosus LLGC-3* | 12.81 | -0.51 | 14.05 | -0.39 | 18.61 | 0.06 | 9.27 | -0.85 | 35.47 | 1.69 |
| *Ananas comosus LLGG* | 98.26 | -0.70 | 86.39 | -0.85 | 134.31 | -0.23 | 161.53 | 0.13 | 278.20 | 1.65 |
| *Phoenix dactylifera LLGE-1* | 729.75 | 1.15 | 452.45 | -0.60 | NA | NA | NA | NA | 157.34 | -0.55 |
| *Phoenix dactylifera LLGE-2* | 0.24 | 0.99 | 33.06 | 0.02 | NA | NA | NA | NA | 16.26 | -1.01 |
| *Phoenix dactylifera LLGH* | 771.78 | -0.99 | 10.26 | 1.01 | NA | NA | NA | NA | 32.53 | -0.02 |
| Basal angiosperm | | | | | | | | | | |
| *Amborella trichopoda LLG* | 768.15 | 1.48 | 171.18 | -0.24 | 42.53 | -0.61 | NA | NA | 39.50 | -0.62 |

Avg. TPM, Average Transcripts Per Million
z-score, standard score after z-transformation of expression values, with each unit representing 1 standard deviation from the mean
Male expression domain included expression in the following tissues: anther, tassel, microspores, pollen grains, pollen tubes, and sperm.
Female expression domain included expression in the following tissues: pistil, stigma, ovary, carpel receptacle, ear, ovule, egg cell, and egg apparatus
RBF, Reproductive before fertilization expression domain included expression in the following tissues: flower, flower buds, ear primordia, immature ears, panicle, and tepals

RAF, Reproductive after fertilization expression domain included expression in the following tissues: zygote, endosperm, nucellus, embryo, seeds, kernel, silique, fruit tissues days after pollination, and pericarp
Veg, Vegetative tissues expression domain included expression in the following tissues: roots, leaves, stems, shoots, meristems, and seedlings
NA, data not available
Gene IDs of Genes listed here can be found in Dataset S1

**Table S5.** List of primers used in this study.

| Primer Number | Primer Description | Sequence (5' – 3') | Template | Primer Set | Expected Length (bp) |
|---|---|---|---|---|---|
| ***pLLG1::LRE-HA*** | | | | | |
| HW1549 | *pLLG1*-FWD1 | AGCGGCCGCGAGGGAGGGTGCTTGAGGTC | *Col-0* gDNA | HW1549 + HW1505 | 2011 |
| HW1505 | *pLLG1*-REV1 | CGGATCCGGTTCTTTGTTGGTTACAGGAG | | | |
| HW1342 | LRE N-terminus FWD 2 | CGGATCCATGGAGCTGATATTATTATTCTTC | *Col-0* gDNA | HW1342 + HW1345 | 60 |
| HW1345 | LRE-GPI-Signal-REV2 | CAGATCTTGAAGAGGAGAGAGATACCAA | | | |
| HW20 | HA-FWD3 | CGGATCCATGGTCTACCCTTATGACGTGCCTG | HA-epitope fragment | HW20 + HW2923 | 80 |
| HW2923 | HA-REV3 | CAGATCTGGCTGCGTAGTCAGGCAC | | | |
| HW1344 | LRE-mature-FWD 4 | CAGATCTAGTTCCATATCGGATGGTGTG | *Col-0* gDNA | HW1344 + HW1343 | 438 |
| HW1343 | LRE-GPI-REV4 | CGTCGACTCAAGTCAACACTAACAAAGC | | | |
| ***pLLG1::LLG3*** | | | | | |
| HW1549 | *pLLG1*-FWD1 | AGCGGCCGCGAGGGAGGGTGCTTGAGGTC | *Col-0* gDNA | HW1549 + HW1505 | 2011 |

| | | | | | |
|---|---|---|---|---|---|
| HW1505 | *pLLG1*-REV1 | CGGATCCGGTTCTTTGTTGGTTACAGGAG | | | |
| HW1890 | *LLG3-cDNA*-FWD | ATGAAGATTACTCATCATTGTTTG | *Col-0* gDNA | HW1890 + HW1891 | 483 |
| HW1891 | *LLG3-cDNA*-REV | TTAGAAGAGGTGAAACAAGATGGA | | | |
| ***pLRE::GUS*** | | | | | |
| 1701 | *pLRE::GUS* FWD Primer 1 | AGGCGGCCGCACTAGTATCTGTGAGTCATCCTTTCGAGGAAATC | *Col-0* gDNA | 1701 + 2195 | 984 |
| 2195 | *pLRE::GUS* REV Primer 1 | GCGGCCGCGAAATTGTTGTTAAAGAAGCTTGTAAC | | | |
| 2196 | *pLRE::GUS* FWD Primer 2 | CAATTTCGCGGCCGCATGTTACGTCCTGTAGAAACC | *pLLG1::GUS* in *Col-0* gDNA | 2196 + 2197 | 1849 |
| 2197 | *pLRE::GUS* REV Primer 2 | AGCGTACCGGACTAGTTCTAGATCATTGTTTGCCTCCCTG | | | |
| 2281 | *pLRE::GUS* FWD Primer 3 | CTAGAACTAGTCCGGTACGCT | *pLRE::LRE-cYFP* plasmid | 2281 + 2282 | 255 |
| 2282 | *pLRE::GUS* REV Primer 3 | AGCTGGGTCGGCGCGCCGATCTGGATTTTAGTACTGGATTTTG | | | |
| ***pLRE::LLG1-cYFP*** | | | | | |
| 1701 | *pLRE::LRE-cYFP* FWD Primer 1 | AGGCGGCCGCACTAGTATCTGTGAGTCATCCTTTCGAGGAAATC | *Col-0* gDNA | 1701 + 1958 | 975 |
| 1958 | *pLRE* REV Primer 1 | GAAATTGTTGTTAAAGAAGCTTGTAAC | | | |

| | | | | | |
|---|---|---|---|---|---|
| 1959 | *pLRE::LLG1-cYFP* FWD Primer 2 | GCTTCTTTAACAACAATTTCATGGAGCTCCTCTCTAGAGC | *Col-0* gDNA | 1959 + 1960 | 994 |
| 1960 | *pLRE::LLG1-cYFP* REV Primer 2 | ACCTCCAGGCCGGCCTACCTCTGCTGATGTC | | | |
| 1961 | *pLRE::LLG1-cYFP* FWD Primer 3 | CAGAGACATCAGCAGAGGTAGGCCGGCCTG | *pLRE::LRE-cYFP* plasmid | 1961 + 1962 | 792 |
| 1962 | *pLRE::LLG1-cYFP* REV Primer 3 | CAGAAGTTTCAGGTGGTAATTGTGACGATCGCTTGTACAGCTC | | | |
| 1963 | *pLRE::LLG1-cYFP* FWD Primer 4 | CACCTGAAACTTCTGCTGAAGTTAACGCAGCAACTACCTCG | *Col-0* gDNA | 1963 + 1964 | 108 |
| 1964 | *pLRE::LLG1-cYFP* REV Primer 4 | TGATTGATCAGAACAACTTAACAAAAACCAAAAGAG | | | |
| 1965 | *pLRE::LLG1-cYFP* FWD Primer 5 | GTTGTTCTGATCAATCAAAGGAAATTGAAAGAGCCA | *pLRE::LRE-cYFP* plasmid | 1965 + 1702 | 158 |
| 1702 | *pLRE::LRE-cYFP* REV Primer 2 | AGCTGGGTCGGCGCGCCGGAGGTCAAGTATTCTTTACACTTGGACACT | | | |
| ***RT-PCR* of *Cleome violacea*** | | | | | |
| 2468 | *Cleome LRE/LLG1* FWD Primer 1 | ATGGAGCTCAAAAGCTTCTCTAG | *Cleome violacea* gDNA | 2468 + 2469 | cDNA: 504 |
| 2469 | *Cleome LRE/LLG1* REV Primer 1 | TCAGAACAACTTGATCAAGACCAAGAA | | | |

| | | | | | |
|---|---|---|---|---|---|
| 2466 | *Cleome ACTIN2* FWD Primer 1 | ATGGCCGAGGAAGCTGATA | *Cleome violacea* gDNA | 2466 + 2467 | cDNA: 1134 |
| 2467 | *Cleome ACTIN2* REV Primer 1 | TTAGAAGCATTTTCTGTGGACGATG | | | |
| ***pLLG1::Clevi-LRE/LLG1-cYFP-3'UTR-LRE*** | | | | | |
| 2301 | *pLLG1::Clevi-LRE/LLG1-cYFP* FWD Primer 1 | GCAGGCGGCCGCACTAGTCGAGGGAGGGTGCTTGA | *Col-0* gDNA | 2301 + 2302 | 2044 |
| 2302 | *pLLG1::Clevi-LRE/LLG1-cYFP* REV Primer 1 | AGCTTTTGAGCTCCATGGTTCTTTGTTGGTTACAGGAGA | | | |
| 2303 | *pLLG1::Clevi-LRE/LLG1-cYFP* FWD Primer 2 | ATGGAGCTCAAAAGCTTCTCT | *Cleome violacea* gDNA | 2303 + 2290 | 1170 |
| 2290 | *pLLG1::Clevi-LRE/LLG1-cYFP* REV Primer 2 | CACCTCCACCTCCAGGCCGGCCAATGTTCGCTGAGGTT | | | |
| 2291 | *pLLG1::Clevi-LRE/LLG1-cYFP* FWD Primer 3 | CTGGAGGTGGAGGTGGAGCTGTGAGCAAGGGCGAG | *pLRE::LRE-cYFP* plasmid | 2291 + 2292 | 768 |
| 2292 | *pLLG1::Clevi-LRE/LLG1-cYFP* REV Primer 3 | TAGCAGAAGTAGGTGGGAGAGCAGGACACGATCGCTTGTACAGCT | | | |
| 2293 | *pLLG1::Clevi-LRE/LLG1-cYFP* FWD Primer 4 | CACCTACTTCTGCTAATATCAACGCTGCCCATATCC | *Cleome violacea* gDNA | 2293 + 2288 | 101 |

| | | | | | |
|---|---|---|---|---|---|
| 2288 | *pLLG1::Clevi-LRE/LLG1-cYFP* REV Primer 4 | ATTTCCTTTGATTGATCAGAACAACTTGATCAAGA | | | |
| 2289 | *pLLG1::Clevi-LRE/LLG1-cYFP* FWD Primer 5 | TCAATCAAAGGAAATTGAAAGAGCC | *Col-0* gDNA | 2289 + 1702 | 148 |
| 1702 | *pLRE::LRE-cYFP* REV Primer 2 | AGCTGGGTCGGCGCGCCGGAGGTCAAGTATTCTTTACACTTGGACACT | | | |
| ***pLRE::Clevi-LRE/LLG1-cYFP-3'UTR-LRE*** | | | | | |
| 1701 | *pLRE::LRE-cYFP* FWD Primer 1 | AGGCGGCCGCACTAGTATCTGTGAGTCATCCTTTCGAGGAAATC | *Col-0* gDNA | 1701 + 1958 | 975 |
| 1958 | *pLRE* REV Primer 1 | GAAATTGTTGTTAAAGAAGCTTGTAAC | | | |
| 2283 | *pLRE::Clevi-LRE/LLG1-cYFP* FWD Primer 2 | TCTTTAACAACAATTTCATGGAGCTCAAAAGCTTCTC | *Cleome violacea* gDNA | 2283 + 2290 | 1187 |
| 2290 | *pLRE::Clevi-LRE/LLG1-cYFP* REV Primer 2 | CACCTCCACCTCCAGGCCGGCCAATGTTCGCTGAGGTT | | | |
| 2291 | *pLRE::Clevi-LRE/LLG1-cYFP* FWD Primer 3 | CTGGAGGTGGAGGTGGAGCTGTGAGCAAGGGCGAG | *pLRE::LRE-cYFP* plasmid | 2291 + 2292 | 768 |
| 2292 | *pLRE::Clevi-LRE/LLG1-cYFP* REV Primer 3 | TAGCAGAAGTAGGTGGGAGAGCAGGACACGATCGCTTGTACAGCT | | | |

| | | | | | |
|---|---|---|---|---|---|
| 2293 | *pLRE::Clevi-LRE/LLG1-cYFP* FWD Primer 4 | CACCTACTTCTGCTAATATCAACGCTGCCCATATCC | *Cleome violacea* gDNA | 2293 + 2288 | 101 |
| 2288 | *pLRE::Clevi-LRE/LLG1-cYFP* REV Primer 4 | ATTTCCTTTGATTGATCAGAACAACTTGATCAAGA | | | |
| 2289 | *pLRE::Clevi-LRE/LLG1-cYFP* FWD Primer 5 | TCAATCAAAGGAAATTGAAAGAGCC | *Col-0* gDNA | 2289 + 1702 | 148 |
| 1702 | *pLRE::LRE-cYFP* REV Primer 2 | AGCTGGGTCGGCGCGCCGGAGGTCAAGTATTCTTTACACTTGGACACT | | | |
